# Molecular and Circuit Architecture of Social Hierarchy

**DOI:** 10.1101/838664

**Authors:** Adam C. Nelson, Vikrant Kapoor, Eric Vaughn, Jeshurun A. Gnanasegaram, Nimrod D. Rubinstein, Venkatesh N Murthy, Catherine Dulac

## Abstract

Social hierarchy is a common organizational structure of animal groups, in which an individual’s social status generates an adaptive behavioral state that facilitates interactions with other group members. Although generally stable, the social status of an animal can change, underscoring the plasticity of the underlying neural circuits. Here, we uncover the molecular and biophysical properties of a cortico-thalamic circuit that supports the emergence of hierarchy in mice. We established a robust behavioral paradigm to explore the establishment of hierarchy among groups of unfamiliar males, and identified the mediodorsal thalamus (MDT) and the caudal part of anterior cingulate area (cACC) as brain areas that are differentially activated between dominants and subordinates. Glutamatergic MDT neurons project to inhibitory parvalbumin interneurons of the cACC, and activity levels of both cell types control competitive performance. Synaptic inputs and excitability of MDT neurons undergo dramatic changes according to the animal social status, and single nucleus sequencing identified increased transcription of the voltage gated ion channel *Trpm3* in dominant mice, leading to enhanced excitatory transmission in the MDT-cACC circuit. Our data suggest a model in which cellular, molecular and biophysical plasticity in a thalamocortical circuit controls the expression of social status.

## INTRODUCTION

Understanding the molecular and circuit mechanisms that underlie flexible behavioral states is a major challenge in neuroscience and neuropsychiatry (Bargmann, 2012; Marder et al., 2014). In a social setting, behavioral states guide how individuals interact with other group members. The social status of an animal is conveyed through dominant (e.g., threatening or aggressive) or submissive (e.g., avoidant or defensive) postures and actions (Frans B. M. de Waal, 1986). These displays are likely adaptive to reduce conflict among individuals with uncertain relationships and provide stability to future interactions (Smith et al., 2011). Social status exerts a far-reaching influence on the brain, from neural activity (Zink et al., 2008) to mental health (Sapolsky, 2005). Although generally stable, social status can change based on alterations in physiological and environmental conditions. Further, hierarchy-dependent interactions require social competence, or the ability to flexibly modulate behavior according to the current social context (Taborsky and Oliveira, 2012).

Processes of hierarchy formation are shared across many species of social animals (Wilson, 1975). In particular, pairwise interactions are the fundamental unit of social hierarchy, which, through iterative steps, lead to orderly networks at the group level (Guhl, 1956; Shizuka and McDonald, 2015). Social hierarchy behavior often emerges spontaneously at a young age, and becomes more structured as individuals prevail or fail in competition over mating and resources (Hawley, 1999). Although more commonly studied in males, females also display social hierarchies (Stockley and Bro-Jørgensen, 2011). Depending on the species, establishment and maintenance of social rank is regulated by multimodal cues, including social signals, memories of previous encounters, and circulating hormones and neuromodulators (Ghazanfar and Santos, 2004). Whereas earlier research documented associations between social status and circulating stress and sex hormones (Creel, 2001; Creel et al., 2012), far less is known about how specific neural circuits are modulated to control flexibility in social hierarchy behavior.

Humans and other primates have an exquisite ability to recognize and enact social hierarchy, and associations with distinctive patterns of brain activity have been identified. FMRI studies of macaques found that activity in cortical areas (temporal sulcus and dorsal and prefrontal cortex) covaried with social status and social network size (Noonan et al., 2014). In experiments where humans participated in direct competitions, fMRI identified distinct areas activated during the participation in hierarchy (Qu et al., 2017). In particular, neural activity in the rostral medial prefrontal cortex (mPFC) matched the dominance status of opponents (Ligneul et al., 2016), while viewing superior individuals engaged the dorsolateral PFC (Zink et al., 2008).

Male and female house mice naturally form hierarchies (Bronson, 1979; Williamson et al., 2019), providing a valuable model system in which underlying mechanistic and circuit-level questions can be directly addressed using molecular and genetic tools. The neural substrates of social rank in males were recently explored using a combination of electrophysiology, virus-based manipulation of neural activity, and behavioral assays such as the tube test, which readily monitors win-or-lose pairwise interactions (Wang et al., 2011; Zhou et al., 2017). In this paradigm groups of co-housed mice spontaneously displayed stable, transitive hierarchies revealed by tube tests and supported by other behavioral assays. Investigation of the mPFC in these established hierarchies revealed greater synaptic strength in dominant vs. subordinate mice in the prelimbic (PL) area of the mPFC. Moreover, constitutive potentiation or depression of α-amino-3-hydroxy-5-methyl-4-isoxazolepropionic acid (AMPA) receptor-mediated synaptic transmission was sufficient to increase or decrease social rank in the tube test (Wang et al., 2011). Subsequent studies of the mediodorsal thalamus (MDT)–showed that synaptic strengthening or weakening of projections of MDT to PL in established hierarchies caused increases and decreases in social rank, respectively (Zhou et al., 2017). Together, these findings showed that plasticity in synaptic transmission underlies aspects of social dominance behavior.

Despite evidence linking plasticity in the mPFC and thalamus with the maintenance of hierarchy in mice, we still lack a brain-wide perspective of the regions and cell-types associated with the progressive establishment of hierarchy among naïve mice. In addition, how the molecular and biophysical properties of functional circuits are modified to control social status remains unclear. Here, we addressed these questions and identified a forebrain-MDT-cACC neural circuit in which molecular and synaptic modulation of activity translates into changes in competitive performance. We showed that glutamatergic MDT neurons and parvalbumin-positive interneurons of the caudal part of the anterior cingulate cortex (cACC) are two interconnected cell types whose activity levels control competitive performance. Using a combination of electrophysiology and single nucleus RNA sequencing, we uncovered the biophysical and molecular underpinnings of the increase in excitability observed in defined MDT cell populations and in MDT-cACC connectivity of high rank males, as well as distinct patterns of synaptic plasticity in inhibitory and excitatory forebrain and cortical inputs to MDT. Together, these data identify molecular, cellular, and physiological mechanisms underlying the emergence of social rank in a critical brain circuit.

## RESULTS

### Emergence of social hierarchy in mice

Recent studies of social hierarchy in mice have used cohorts of co-housed male mice, such that a fraction of the tested animals spontaneously display stable hierarchical relationships in tube test and other dominance-based assays (Wang et al., 2011; Zhou et al., 2017). Here, we aimed to uncover the neural mechanisms underlying the emergence of stable hierarchy in a naïve cohort. For this purpose, we sought to develop a robust behavioral paradigm in which hierarchical relationships among single-housed, unfamiliar adult male mice are established and monitored. Cohorts of four single-housed adult C57BL/6J males (∼ eight weeks old) were introduced to each other through pairwise interactions in two distinct round robin assays: resident-intruder (RI) and tube tests (TT), each performed daily over approximately one week (Fig. 1A & B). In resident-intruder assays, agonistic behavior (i.e. defensive or aggressive) is elicited by introducing an intruder into the home cage of the resident (Miczek et al., 2001). In tube test assays, two mice interact in a tube large enough to allow head-to-head interactions— but not aggressive fighting—until one mouse, the loser, exits the tube (Fan et al., 2019). Iterative tube test tournaments provide clear binary outcomes for each trial and have been shown to provide a readout of hierarchical relationships supported by the outcomes of other dominance-subordinate assays tested (Wang et al., 2011).

**Figure 1.**
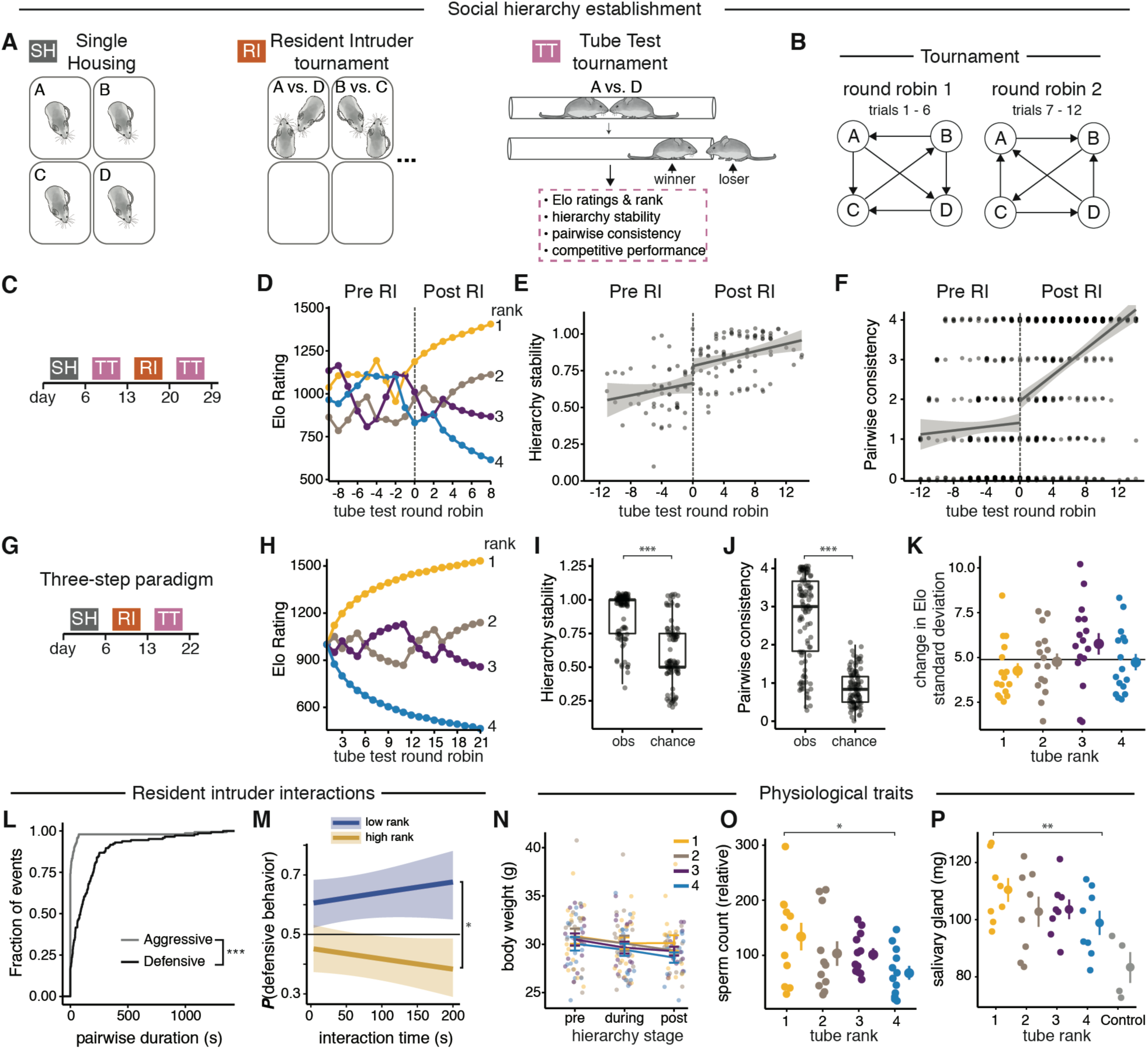
Determinants of hierarchy formation. **(A - B)** Visualization of behavioral assays and tournament structure. **(A)** Left: single housing (SH) prior to, and throughout, experimental period. Middle: resident-intruder (RI) tournament (trials repeated daily for at least one week). Right: tube test (TT) tournament (trials repeated daily for at least one week). **(B)** Schematic of tournament structure. Round robin 1: each pairwise combination of animal encounters. Round robin 2: same as round robin 1 but in reverse order. **(C-F)** Social interactions during resident-intruder facilitate hierarchy displayed in tube tests. N = 8 groups of 4 mice. **(C)** Schematic of a four-step experimental paradigm. **(D)** Tube test Elo ratings and ordinal ranks of an example group of four mice before and after a resident-intruder tournament Tube ranks are color coded as follows: Rank-1 (gold), Rank-2 (light-brown), Rank-3 (purple), Rank-4 (blue). Hierarchy stability **(E)** and pairwise consistency **(F)** across repeated tube test tournaments before and after a resident intruder tournament. Dots show the stability index per round robin per group in (E) and the consistency value per pairwise interaction per group in (F). Lines show the predicted values of the linear model ± s.e.m. **(G)** Three-step paradigm for the emergence of social hierarchy among unfamiliar mice. **(H)** Elo ratings and ordinal ranks of example group of four mice across 21 round robins. Tube test hierarchy stability **(I)** and pairwise consistency **(J)** of observed groups compared to random outcomes, box plots show median and first and third quartiles, LMER, N = 12 groups of 4 mice. **(K)** Standard deviation in delta Elo scores by tube test rank, LMER, N = 15 groups of 4 mice. Horizontal line denotes mean standard deviation. **(L)** Cumulative distribution plot showing the fraction of aggressive vs. defensive events displayed during resident-intruder interactions. N = 12 groups of 4 mice. **(M)** Probability of defensive behavior as a function of resident-intruder interaction time for high rank and low rank mice. Binomial logistic regression ± s.e.m. **(N-P)** Physiological correlates of social rank. **(N)** Body weight during the three-step hierarchy formation paradigm according to rank. N = 18 groups of 4 mice (72 mice). Quantification of relative sperm count **(O)** and salivary gland mass **(P)** in relation to final tube rank or individually housed control males, LMER with Tukey post hoc test, mean ± s.e.m. N = 4 groups of 4 mice plus 2 control mice (18 mice).

Hierarchical relationships in tube tests were scored with Elo ratings, which continuously update player ratings based on whether performance is better or worse than expected from previous contests (Elo, 1978; Neumann et al., 2011). With Elo ratings, expected outcomes lead to smaller point changes than unexpected outcomes (SOM Methods). Thus, probabilistic models of dominance like Elo ratings provide an advantage over strict ordinal ranks because they can accommodate players tied for number of wins, quantify the magnitude of dominance between ranks, and are amenable to parametric statistics controlling for multiple response variables (Boyd and Silk, 1983). Where appropriate, we used Linear Mixed Effect Models (LMERs) (Bates et al., 2015) to identify the correlates of social rank, which allowed us to control for between-group variation (i.e., “group effects”) and batch effects (e.g., due to batch processing). In addition, we monitored the stability of hierarchies within individual groups using two metrics. First, we determined a hierarchy stability index measured as the ratio of rank changes per individual in the group over a given time period, such that a stability index of one indicates no rank changes, while an index of zero indicates rank changes for all four animals. Second, we measured a pairwise consistency index, which counts consistent interactions in a sliding window for each pair of mice in the group. The index resets to zero each time the outcome changes, and reaches a maximum equivalent to the number of mice in the group (e.g., four in a group of four).

When groups of four mice were assessed in tube test tournaments prior to any mutual social contact through resident-intruder (RI) interactions, the groups displayed relatively low hierarchy stability (i.e., more rank changes) and low pairwise consistency (i.e., less consistent outcomes for each pairwise interaction) (Fig. 1C-F Pre RI). By contrast, an initial exposure of mice to territorial interactions through resident-intruder round robin assays readily promoted the establishment of hierarchy, as revealed by tube tests (Fig. 1C-F Post RI). These observations led us to evaluate hierarchy establishment in a three-step paradigm: (1) single housing, (2) resident-intruder tournament, and (3) iterated tube test tournaments, with each step lasting approximately one week (Fig. 1G). Under this paradigm, groups of mice exhibited clear differences in social rank according to Elo rating (Fig. 1H), as well as more stable hierarchies (Fig. 1I) and greater pairwise consistency (Fig. 1J) than would be expected by chance. To address whether there were differences in the magnitude of dominance between ranks, we calculated the variance in the competitive performance of each animal, where competitive performance is defined as the per-trial change in Elo rating. We found that rank-1 mice had below-average, and rank-3 mice had above-average variance in competitive performance (Fig, 1K, Bartlett’s test, N = 16 groups, P < 0.001), suggesting that rank-1 mice are less likely, and rank-3 mice are more likely, to deviate from their rank.

Similar to groups of four mice, groups of three mice subject to this three-step paradigm showed high levels of hierarchy stability and pairwise consistency. In addition, rank-1 mice showed lower variance in competitive performance compared to rank-2 and rank-3 mice (SOM Fig. 1A-E). The establishment of social hierarchy using this three-step paradigm (Fig. 1G) applied to groups of three or four initially unfamiliar mice served as the basis for all subsequent experiments and manipulations.

We next assessed how agonistic interactions during resident-intruder trials predicted outcomes in the tube test. We found that defensive behaviors (i.e., fleeing or avoiding the other mouse) were present in every trial and were displayed far more frequently than aggressive (i.e., attack) behaviors (Fig. 1L). We then used logistic regression to model the probability of displaying agonistic behavior during resident-intruder assays and found that mice achieving high ranks (rank-1 and rank-2) were less likely to be defensive than those achieving low ranks (rank-3 and rank-4) (P < 0.05, Fig. 1M), whereas the probability of attack behavior was not associated with rank (p > 0.05, SOM Fig. 1F). Next, we found that the relative duration of defensive behavior (calculated as A/(A+B) for mouse A competing against mouse B) was a better predictor of tube rank than the absolute duration of defensive behavior, as determined by the percent of variance explained (R^2^ values for defensive behavior: relative duration = 0.22, absolute duration = 0.02, SOM Fig. 1G). By contrast, both the relative duration of attacks and the absolute duration of attacks were poor predictors of tube rank (R^2^ values for attacks: relative duration = 0.08, absolute duration = 0.09). These results suggest that mice modulate their defensive responses based on the status of the competitor they are interacting with, and that increased defensive behavior in low rank rather than aggressive behavior in high rank marks hierarchy formation.

Together, these data validate our newly developed behavior paradigm as a rigorous tool to capture hierarchy formation, such that experience-dependent processes lead to stable social interactions, and defensive behavior is associated with social rank.

### Physiological correlates of social rank

We next investigated physiological traits and social cues associated with hierarchy in groups of four. Changes in body weight have been associated with rank establishment in some species (Huchard et al., 2016). We found that although mice lost weight during hierarchy formation, social rank was not correlated with weight before, during or after establishment of hierarchy (LMER, Tukey post hoc comparisons, all p > 0.05, Fig. 1N). Testosterone levels have been associated with the display of aggression but evidence has been so far contradictory (Nelson and Trainor, 2007), likely due to the rapid and high fluctuations in testosterone levels, and the low specificity of most detection methods (Rosner et al., 2007; Van Uytfanghe et al., 2004). We therefore measured traits known to be androgen dependent (Sawada and Noumura, 1991; Smith and Walker, 2014) as a readout of testosterone levels, and found that sperm count and salivary gland mass were higher in rank-1 compared to rank-4 males (Fig. 1O,P). Salivary glands produce aggression-promoting pheromones (Taha et al., 2009), raising the possibility that winners express more androgen-dependent pheromones in saliva (Gröschl, 2009).

To investigate the role of chemosensory cues, we measured competitive performance during tube test interactions when saliva, urine, or both were swabbed between rank-1 and rank-4 mice within a group. Saliva had no effect on tube test interactions. In contrast, urine from rank-1 males transiently increased competitive performance of rank-4 males in 66% of trials. Rank-4 mice returned to the bottom rank on subsequent trials in which they were not swabbed (SOM Fig. 1H, I). Transferring urine from rank-4 to rank-1 had no effect. To investigate whether other sensory cues were associated with hierarchical relationships, we measured ultrasonic vocalizations (USVs) and found that USVs were not emitted during tube tests (SOM Fig. 1J). Together, these results suggest that social rank is associated with androgen-dependent traits and confirmed previous observations demonstrating that urinary cues of dominant males signal high rank (Hurst et al., 2001; Nelson et al., 2015).

### Neural activation patterns during tube test tournaments

To identify brain areas involved with social hierarchy, we compared the number of cells expressing the immediate early gene *Cfos* in rank-1 *vs.* rank-4 males following tube test tournaments (Fig. 2A), as well as in individually housed controls passed through the tube without social interaction (SOM Fig. 2A). Using fluorescent in situ hybridization (FISH) and an automated image analysis pipeline (SOM methods) we analyzed 25 brain regions associated with primate social hierarchy (Chiao, 2010) or seen as activated in pilot experiments.

**Figure 2.**
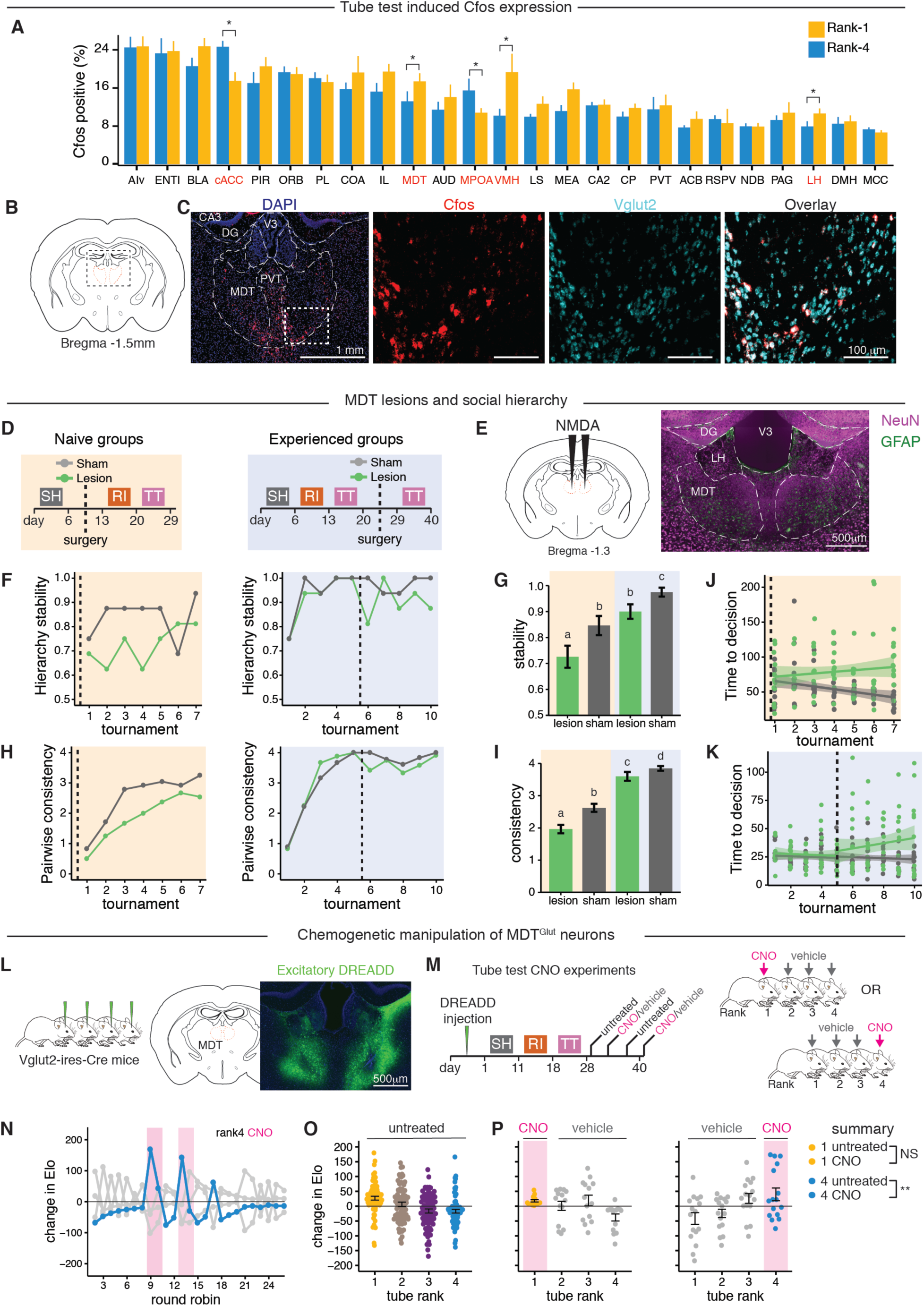
Social rank-dependent brain activity and MDT requirement in emergence of social hierarchy. **(A-C)** Identity and activity of brain areas associated with rank in the tube test. **(A)** Quantification of *Cfos* expression in 25 brain regions from rank-1 and rank-4 males, mean ± s.e.m. LMER with Tukey post hoc test. N = 4 social groups, * p < 0.05. **(B)** Schematic of the MDT. **(C)** MDT-glut neurons are *Cfos*+ during tube test interactions. Schematic shows anatomical location of MDT FISH. Also shown are DAPI+ cells (including regions proximal to MDT), *Cfos*+ cells, *Vglut2*+ cells, and *Cfos*-*Vglut2* overlap. **(D-K)** MDT is required for emergence of stable hierarchy. **(D)** Schematic of experimental timeline. Lesions were performed in groups of four animals prior to hierarchy establishment (Naïve, light orange panels), or afterwards (Experienced, light blue panels). Sham groups received saline injections. **(E)** Schematic of NMDA injections to the MDT (left) with NeuN and GFAP staining of a lesioned MDT (right). **(F-K)** Effects of MDT lesion on hierarchy dynamics. Mean hierarchy stability **(F)** and consistency **(H)** during tube test tournaments in naïve (left) and experienced (right) groups compared to sham controls. Summary of stability **(G)** and consistency **(I)** measurements; Wilcoxon rank sum test, bars not connected by same letter are significantly different (p<0.05). Tube test decision timing in naïve **(J)** and experienced **(K)** groups compared to sham controls; fitted lines are the predicted values of linear model. Mean ± s.e.m, N = 8 social groups. **(L)** Viral mediated conditional expression of DREADDs in all four males within a group. Anatomical location of MDT (right) and expression of excitatory DREADD in Vglut2-ires-Cre mouse. **(M)** Experimental timeline. After two weeks of recovery and viral expression, groups established hierarchy. CNO was delivered during tournaments according to a repeated on-off schedule: during “on” tournaments, either rank-1 or rank-4 males were treated with CNO, while the remaining three group members received vehicle. During intervening “off” tournaments all mice were untreated. (**N-P)** Effect of excitation of MDT^Glut^ on social rank, means ± s.e.m, LMER, N = 4 groups. ** p<0.01. ***p<0.001.

To best identify possible links between *Cfos* expression and social rank we used LMER models to control for group effects and batch effects of histology processing. We found that Rank-1 and rank-4 mice exhibited largely similar patterns of *Cfos* expression, both in terms of the identity of brain areas expressing *Cfos* and in the number of *Cfos*-positive cells within those areas. However, we uncovered a significant effect of social rank on the number of *Cfos*-positive cells in five brain areas, including areas previously associated with agonistic social behavior (Fig. 2A). Compared to rank-4, rank-1 males had increased activity in the ventromedial hypothalamus, an area required for aggression (Lin et al., 2011), and in the lateral habenula, a brain structure associated with aggression reward (Golden et al., 2016) and resolution of aggressive conflicts (Chou et al., 2016). Rank-1 males had decreased activity in the anterior cingulate area, including the caudal part of anterior cingulate cortex (cACC) proximal to the genu of corpus callosum, a region involved in the expression of conditioned fear (Courtin et al., 2013; Frankland et al., 2004) and emotional conflicts (Davis et al., 2005), and in the medial preoptic area (MPOA), a conserved circuit node in a variety of social behaviors including aggression (Moffitt et al., 2018; Wu et al., 2014) and social reward (McHenry et al., 2017). Rank-1 males had also increased activity in the mediodorsal thalamus (MDT), a finding consistent with the previous observation that ectopic activation of MDT projections to the mPFC affected social rank (Zhou et al., 2017).

To explore potential functional connectivity among the 25 brain regions we evaluated differences in pairwise correlations in *Cfos* expression across brain regions according to social rank (SOM Fig. 2B). We uncovered nine pairwise correlations that were significantly different between rank-1 and rank-4, which, intriguingly, were all associated with either MDT (4/9) or cACC (5/9) (SOM Fig. 2B, right). Social rank affected the correlation of *Cfos* expression in the MDT with three cortical areas (prelimbic area (PL), orbitofrontal cortex (OFC), and cACC as well as with the hippocampus CA2. Social rank affected the correlation of *Cfos* expression in the cACC with the MDT, lateral habenula (LH), nucleus of the diagonal band (NDB), lateral septum (LS), and nucleus accumbens (ACB) (SOM Fig. 2B). These activity-based correlations are supported by previous neuroanatomical tracing studies: MDT directly connects with ACC, PL, and OFC (Kuroda et al., 1998), and ACC directly connects with NDB, LS, and MDT (Fillinger et al., 2017, 2018). Moreover, the MDT-PL circuit is an established node in the control of social hierarchy (Zhou et al., 2017), CA2 is required for social memory processing (Hitti and Siegelbaum, 2014), LS is associated with anxiety and aggression (Beiderbeck et al., 2007), and LH is associated with aggression (Golden et al., 2016).

The functional organization of the MDT is characterized by excitatory thalamocortical projections and a lack of local connections (Halassa and Kastner, 2017; Kuroda et al., 1992). Accordingly, we found that *Cfos*-positive cells in rank-1 and rank-4 were uniformly *Slc17a6-* positive (i.e., VGLUT2-positive) glutamatergic neurons (MDT^Glut^, Fig. 2B, C & SOM Fig. 2C). Together, these results indicate that social rank is associated with distinct patterns of neural activity, particularly in MDT^Glut^ and cACC neurons.

### MDT facilitates the establishment of hierarchy

In our behavioral paradigm, stable hierarchy emerges after multiple territorial interactions (Fig. 1C-F), suggesting the existence of underlying experience-dependent processes. Recent studies have suggested that mutual connectivity between MDT and cortex guides elementary cognitive processing, adaptive decision making, and flexible behavior, particularly in the context of multimodal signal processing (Chakraborty et al., 2016; Parnaudeau et al., 2015; Schmitt et al., 2017) and that plasticity of MDT to PFC inputs mediate changes in social dominance (Franklin et al., 2017; Zhou et al., 2017). To further address the role of MDT in the formation and maintenance of social hierarchy, we lesioned this structure either before or after hierarchy formation using targeted NMDA (N-methyl-D-aspartate) injections, which eliminates cell bodies but not fibers of passage (Maren et al., 1997) (Fig. 2D, E).

In “naïve” cohorts, lesions were delivered to all four mice in a social group prior to hierarchy formation; in “experienced” cohorts, lesions were delivered to all four mice after hierarchy establishment. Animals receiving sham surgeries with vehicle injections served as controls for both cohorts (Fig. 2D). Lesions did not affect aggressive behavior during resident-intruder assays tested in naïve cohorts (SOM Fig. 2D). Analysis of post-operative social dominance dynamics showed that animals receiving lesions before hierarchy formation displayed more variable Elo ratings (SOM Fig. 2F). Accordingly, compared to sham control groups, naïve lesion groups exhibited robust decreases in hierarchy stability and pairwise consistency, whereas experienced lesion groups exhibited only modest decreases (Fig. 2F-I and SOM. Fig.2E-F). Lesions also resulted in longer post-operative tube test decision timing in naïve and experienced groups (Fig. 2J-K).

MDT receives inputs from olfactory cortical areas (Courtiol and Wilson, 2015; Price, and Slotnick, 1983) including the OFC, an area associated with odor reward value (Schoenbaum and Eichenbaum, 1995). We tested whether MDT was required for the odor response to rank-1 urinary cues observed in swabbing experiments (SOM Fig.1H-I). As expected, baseline competitive performance in the untreated condition (i.e., no urine swab) was positively correlated with final tube rank (SOM Fig. 2G & I), and, in sham groups, rank-4 males swabbed with rank-1 urine had increased performance (SOM Fig. 2H). In contrast, groups with MDT lesions failed to show the same response: rank-4 performance was unaffected by the urine swab (SOM Fig. 2J), suggesting that MDT is required for executing appropriate hierarchy-dependent responses to urinary olfactory cues.

Next, we investigated whether MDT more widely contributes to sociability or social recognition using a three-chamber sociability assay and found no differences between the behavior of lesion and sham animals (SOM Fig. 2K-L). Taken together these results suggest that, although MDT is not required for sociability nor social recognition, it is required for establishing a stable hierarchy and consistent interactions.

### MDT activation increases competitive performance

*Cfos* expression in the MDT was associated with winning in the tube test, prompting us to examine whether chemogenetic manipulation of MDT^Glut^ neurons was sufficient to affect competitive performance. Separate cohorts of four mice were bilaterally infected with conditional AAVs encoding either inhibitory or excitatory DREADDs in MDT^Glu^ (Fig. 2L) and allowed to establish hierarchies. DREADDs are engineered receptors activated by the cognate ligand CNO (Alexander et al., 2009; Armbruster et al., 2007). CNO was delivered according to a repeated on-off schedule: during “on” tournaments, either rank-1 or rank-4 males were treated with CNO while the remaining three group members received vehicle (Fig. 2M). During intervening “off” tournaments all mice were untreated. In initial assays, we tested CNO injections in rank-1 and rank-4 mice in the absence of DREADDs and found that CNO alone did not affect the established hierarchy in control groups of C57/B6 mice (LMER, P >0.05) (SOM Fig 3A).

**Figure 3.**
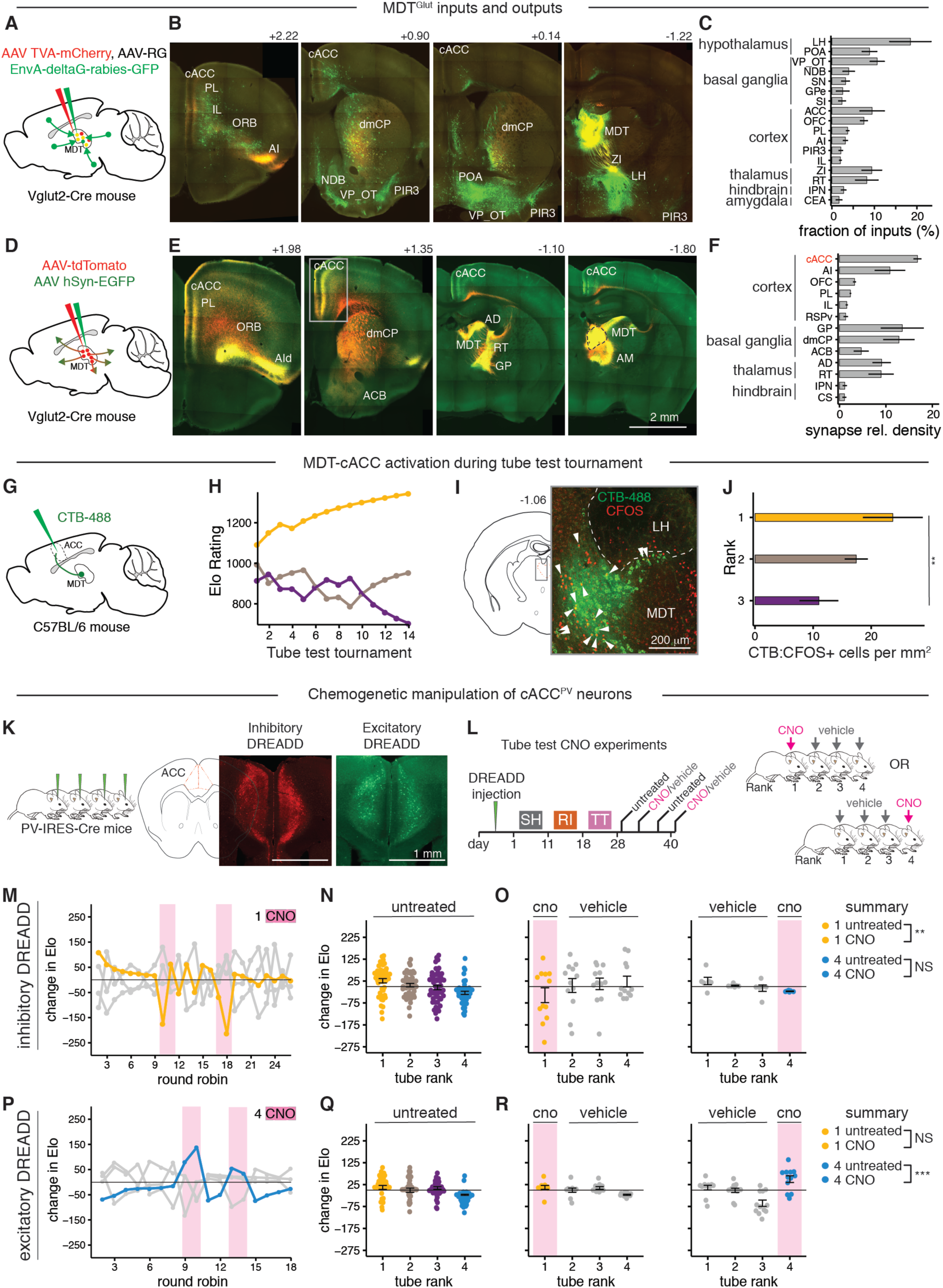
Inputs and outputs of MDT^Glut^ neurons. **(A-C)** Monosynaptic retrograde tracing from MDT^Glut^ neurons. **(A)** Tracing scheme. **(B)** Input areas displaying rabies-positive neurons in control male. **(C)** Input fractions to MDT^Glut^ neurons (percent of total counted inputs), mean ± s.e.m, N = 3 mice. **(D-F)** Anterograde tracing of MDT^Glut^ neurons. **(D)** Tracing scheme. **(E)** Afferent projection synapses (green) and fibers (red) in a control male. **(F)** Relative density of synapses; mean ± s.e.m, N = 3 mice. **(G-J)** Neural activity of MDT-cACC projections during competition. **(G)** Tracing scheme. **(H)** Example Elo ratings from a group of three prior to CFOS labeling. **(I)** Labeling of CTB (green), CFOS IHC (red), and co-labeling (white arrows) from a rank-1 mouse. **(J)** Quantification of CTB-CFOS cells by rank, mean ± s.e.m, LMER, N = 12 mice ** p <0.01. **(K)** Viral mediated conditional expression of DREADDs in all four males within a group (left). Anatomical location of cACC (right) and expression of inhibitory (red) and excitatory (green) DREADDs in *PV*-Cre mouse. **(L)** Experimental timeline. After two weeks of recovery and viral expression, groups established hierarchy. CNO was delivered during tournaments according to a repeated on-off schedule: during “on” tournaments, either rank-1 or rank-4 males were treated with CNO, while the remaining three group members received vehicle. During intervening “off” tournaments all mice were untreated. **(M-O)** Effect of inhibition of cACC^PV^ on social rank. **(M)** Individual performance (change in Elo) in a hierarchy where the rank1 male (yellow trace) received CNO (shaded rectangles). Gray lines represent the other three males. **(N)** Relationship between performance and final tube rank in untreated mice. **(O)** Effect of CNO in experiments where either rank-1 (left) or rank-4 (right) received CNO while the other three mice received vehicle. Statistical effects are summarized on the right, means ± s.e.m, LMER, N = 3 groups. **(P-R)** Effect of excitation of cACC^PV^ on social rank. Means ± s.e.m, LMER, N = 3 groups.

In cohorts receiving chemogenetic excitation of MDT^Glut^ neurons, CNO transiently caused a robust competitive performance spike in rank-4 males (Fig. 2N-P) (SOM Video 1 of tube test), but did not affect rank-1 males (Fig. 2P). Chemogenetic inhibition did not affect competitive performance in either rank-1 or rank-4 males (SOM Fig 3B-D). Thus, excitation of MDT is sufficient to increase competitive performance of low rank males while, in agreement with NMDA lesions, suppression of MDT activity after the emergence of hierarchy has no effect on an animal’s ranking.

### Organization of MDT inputs and outputs

Having established a functional role of the MDT in the establishment of social hierarchy, we next sought to define the inputs and outputs of MDT^Glut^ neurons. We assessed the brain-wide inputs to MDT^Glut^ neurons in Vglut2-ires-Cre mice using (G)-deleted rabies-based retrograde trans-synaptic labeling (Wickersham et al., 2007) with helper AAVs (Fig. 3A). Our data show that nearly 20 areas sent mono-synaptic inputs to MDT^Glut^ neurons, including hypothalamic areas associated with arousal, feeding, and pain (e.g., lateral hypothalamus (LH), preoptic area (POA)), basal forebrain areas associated with reward learning and voluntary movement (e.g., NDB, ventral pallidum (VP), substantia innominata (SI)), deep-layer cortical areas associated with executive function (ACC, OFC, PL), areas associated with olfaction (piriform cortex, olfactory tubercle, OFC), and intra-thalamic information processing areas (reticular thalamus and zona incerta) (Fig. 3B-C).

Projections from MDT^Glut^ neurons were visualized using two combined AAVs driving expression of reporter fluorophores in axons (tdTomato) and presynaptic terminals (Synaptophysin-GFP fusion protein) (Fig. 3D). As previously described, layer II/III in cortical areas associated with executive, perceptual and memory functions (e.g., ACC, OFC, PL) received the majority of MDT^Glut^ projections (Delevich et al., 2015; Krettek and Price, 1977). The ACC, from its most rostral (> bregma +2.0) to caudal (< bregma – 2.0) coordinates, received the greatest fraction of inputs (Fig. 3E-F). Additional targets included cortical (retrosplenial and infralimbic cortex) and basal ganglia areas (i.e., globus pallidus, dorsomedial caudate putamen, and nucleus accumbens). The MDT^Glut^ circuit is therefore poised to integrate aspects of learned interactions, emotional valance, innate behavioral responses, and olfactory communication, all functions that are highly relevant to hierarchy formation.

### Activity of MDT-cAAC projections during social competition

The cACC, a major target of MDT^Glut^ projections, showed decreased activity in rank-1 males after tube tests (Fig. 2A). Because rank-1 males showed increased activity in MDT, we hypothesized that MDT may inhibit the cACC. Indeed, reports using viral tracing (Delevich et al., 2015; Lu et al., 2017), optogenetic assisted circuit mapping (Delevich et al., 2015; Ferguson and Gao, 2018; Miller et al., 2017), and neurophysiological studies of decision making (Schmitt et al., 2017) suggest that MDT projections can drive feedforward inhibition of the PFC through connections with PV interneurons (Delevich et al., 2015). We confirmed this connectivity and assessed its relevance to neuronal activity associated with hierarchy with the following experiments. First, we performed retrograde tracing from PV interneurons in the cACC (cACC^PV^, SOM Fig. 3E-F). While the majority presynaptic input of cACC^PV^ were local (∼79% of inputs), we detected input cells in the lateral MDT (∼5%, SOM Fig. 3G), a subdivision of the MDT that was *Cfos*-positive during tube tests (Fig. 2C). Additional input cells were found in three thalamic areas (anteromedial nucleus (∼11%), lateral dorsal nucleus (>1%), and nucleus reuniens (>1%)), as well as the ventral pallidum (∼3%) (SOM Fig. 3C).

We next asked whether MDT-cACC projection activity was associated with social rank. We injected the retrograde tracer CTB into the cACC of each animal in established three-mouse hierarchies (Fig. 3G-H) and measured neural activity using CFOS immunostaining following tube tests (Fig. 3I-J). Rank-1 mice had more double-labeling of CTB-CFOS cells than rank-3 mice in the lateral MDT (Fig. 3I-J). These results suggest that lateral MDT^Glut^-cACC projections are (1) capable of driving feedforward inhibition, and (2) differentially activated according to social rank.

### cACC^PV^ neurons regulate competitive performance in social contests

Based on our findings that social rank is associated with levels of *Cfos* expression in the MDT-cACC circuit and that cACC^PV^ cells are a significant target of activated MDT^VGlut^ cells during hierarchical encounters (Fig. 2A and Fig. 3J), we next tested whether chemogenetic manipulation of cACC^PV^ interneurons was sufficient to control competitive performance and social rank during tube tests (Fig. 3K-L). The ACC is positioned between limbic and cortical brain structures to consolidate emotions and perception, thus shaping amygdala-dependent fear learning and memory (Frankland et al., 2004; Toyoda et al., 2011). In addition, classical conditioning experiments have shown that inhibition of PV interneurons of the mPFC and ACC leads to fear expression (Courtin et al., 2013), a finding perhaps consistent with greater *Cfos* expression in the cACC of lower ranking mice. We therefore predicted that inhibition (or excitation) of PV interneurons would lead to a decrease (or increase) in competitive performance.

Indeed, in cohorts expressing inhibitory DREADD in cACC^PV^ interneurons, CNO caused a pronounced, transient decline in competitive performance among rank-1 males but did not affect the performance of rank-4 males (Fig. 3M-O). By contrast, in cohorts expressing excitatory DREADD in cACC^PV^, CNO caused a pronounced, transient competitive performance spike in rank4 males, but did not affect rank-1 males (Fig. 3P-R). Thus, cACC^PV^ neurons exert a bidirectional control of competitive performance in the tube test, with excitation driving increases, and inhibition driving decreases in performance. Together, these results are consistent with a model in which MDT^Glut^ activation of cACC^PV^ increases performance during competitive social interactions.

### Role of cACC^PV^ and MDT^Glut^ neurons outside of hierarchy

Results from lesion and chemogenetic studies supported the notion that the MDT and cACC are involved in aspects of social hierarchy behavior. To address the specificity of these findings, we next investigated how chemogenetic manipulation of cACC^PV^ and MDT^Glut^ neurons affects anxiety-like behavior and sociability and social recognition. For this purpose, we performed open field and 3 chamber social interaction assays (SOM Fig. 3I-R). There was no effect of rank on open field behaviors (velocity, distance travelled, time spent in the inner chamber, ANOVA, all p values > 0.05), or on social interaction behaviors (sociability and social recognition, all p values > 0.05). Animals of all ranks were therefore pooled together to test the effects of DREADD-based manipulations on behavior in the open field. Similarly, for cACC^PV^ experiments, there was no effect of DREADD construct (p > 0.05), so animals receiving excitatory or inhibitory constructs were also pooled together

In cACC^PV^ activity manipulations, we observed an unexpected baseline effect in absence of CNO injection and according to the type of DREADD receptor expressed: mice in which cACC^PV^ expressed inhibitory DREADD spent less time in the open field and had greater velocity than mice in which cACC^PV^ expressed excitatory DREADD, suggesting higher baseline anxiety (SOM Fig. 3H-J). However, there was no effect of CNO excitation or inhibition of cACC^PV^ neurons on behavior in the open field (SOM Fig. 3H-J). In MDT^Glut^ chemogenetic excitation experiments, CNO caused an increase in time spent in the open field and a marginal decrease in velocity and distance traveled (SOM Fig. 3M-O). DREADD-based manipulations of cACC^PV^ or MDT^Glut^ did not affect sociability or social recognition (SOM Fig. 3 K-L & 3 P-Q). These results imply that activation of MDT^Glut^ is marginally anxiolytic, but neither activity of cACC^PV^ or MDT^Glut^ has any effect on sociability or social recognition.

### Biophysical and molecular characteristics of MDT neurons associated with the establishment of social rank

To address whether the establishment of social rank was associated with persistent changes in intrinsic MDT neuronal properties, we performed whole-cell patch clamp recordings of MDT neurons in acute slices from each individual animal from cohorts of three mice with established hierarchies, as well as individually housed controls, one day or more after the last tube test (Fig. 4A and SOM Fig. 4A-B). We addressed possible differences in intrinsic membrane properties of MDT neurons according to the individual’s social rank by recording from MDT cell bodies in current-clamp mode, and promoted spike firing by injecting variable amounts of current. Across the population, MDT neurons showed a bimodal distribution of peak firing rates, with some exhibiting low, and others high, firing frequencies (Fig. 4B and SOM Fig. 4C-D). MDT cells also showed a bimodal distribution of membrane resistances (Fig. 4B). We verified the bimodality of the data (Hartigans’ dip test, p = 0 for both parameters), and split the MDT neurons into two clusters (k-means clustering). Cluster-1 contained 62% (37/60) of the recorded neurons and was characterized by low-firing frequency and low membrane resistance (membrane resistance <500 MΩ and peak firing rate <65 Hz), while cluster-2 contained 38% (23/60) of the recorded neurons, which displayed high-firing frequency and high membrane resistance (membrane resistance >500 MΩ and peak firing rate >50 Hz) (Fig. 4C). Intriguingly, cells with high-firing rates and high membrane resistance were more likely to originate from rank-1 males, whereas cells with low-firing, low membrane resistance, were more likely to originate from rank-2, rank-3 and control males (Fig. 4C-D). Strikingly, whereas rank-2, rank-3, and control animals had similar and relatively low ratios of cluster-2 (high frequency firing) to cluster-1 (low frequency firing) cells, rank-1 animals had an approximately five-fold higher ratio of high versus low firing rate neurons. This observation suggested that MDT neurons may exhibit two distinct states of excitability according the animal’s social rank.

**Figure 4.**
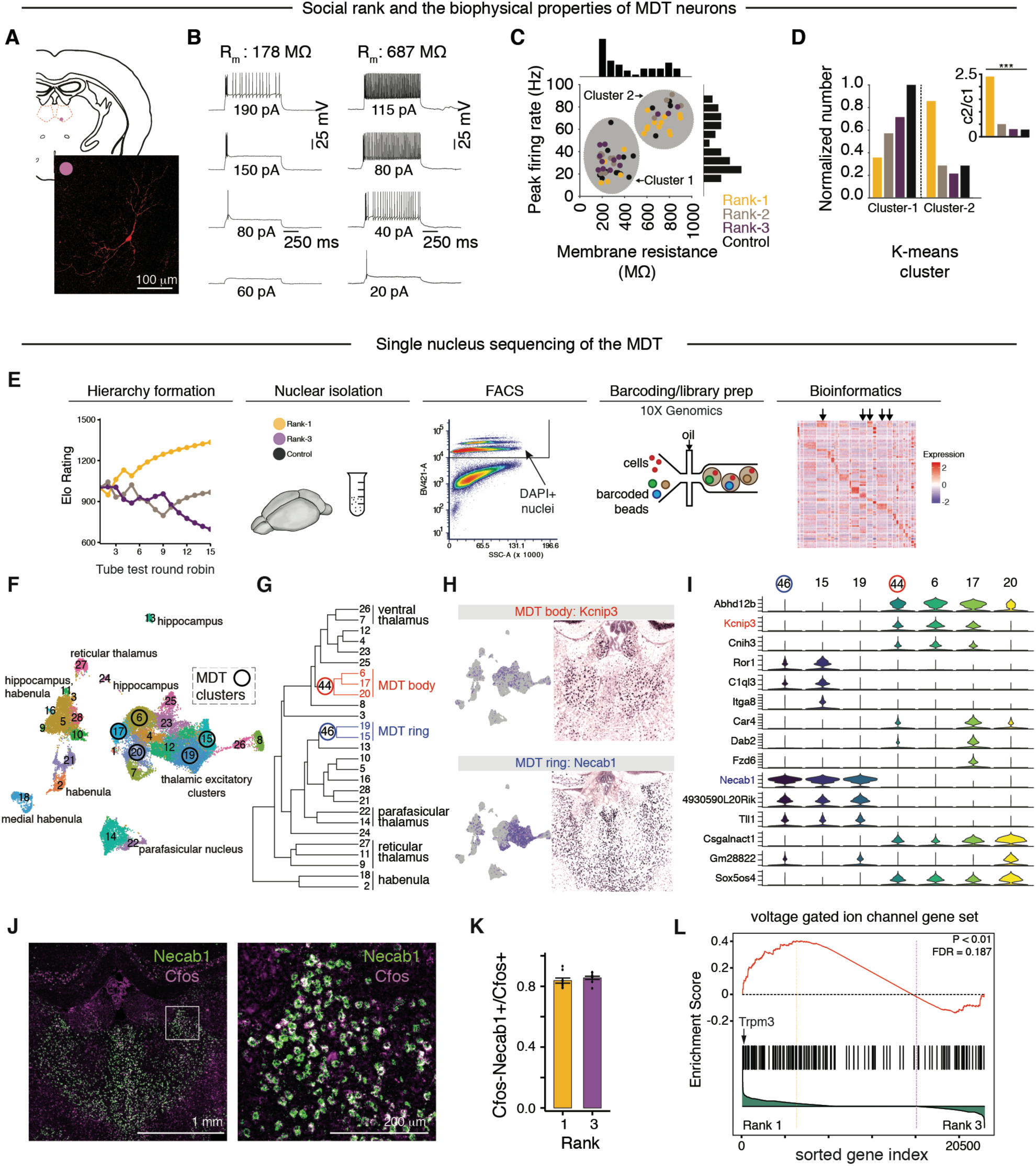
Effects of hierarchy formation on intrinsic properties of MDT neurons. **(A)** Schematic of the location (top) and staining (bottom) of example of recorded cell from MDT. **(B-D)** Whole-cell current clamp recordings of MDT neurons. N = 60 neurons. **(B)** Example traces of Cluster-1 type (Low-firing, low-resistance; left) and Cluster-2 type (high-firing, high-resistance (right) neurons. R_m_, membrane resistance. **(C)** Scatter plot comparing neuronal peak firing rate and membrane resistance according to cluster-type. Hartigans’ dip test confirmed bimodality of data, p = 0 for N=33 neurons in cluster-1 and N=27 neurons in cluster-2. Histograms show bimodality of the data for membrane resistance (top) and for peak firing rate (right). **(D)** Summary depicting effect of social rank on the proportion of cluster-types according to social rank. Inset: ratio of cluster-2/cluster-1 neurons in MDT according to rank,***p<0.001. **(E-I)** Molecular characterization of hierarchy-dependent changes occurring in MDT ring cells. **(E)** Overview of experimental strategy. Stable social hierarchies among groups of three mice were established. Nuclear suspensions were made from dominant (rank-1), subordinate (rank-3) and control mice. DAPI-positive nuclei were isolated by FACS. Single-nucleus libraries were prepared using the 10x Genomics microfluidics platform. Raw reads were processed to obtain gene expression matrices, and clusters (and markers) were identified using a smart local moving algorithm to cluster on shared nearest neighbors in PCA-space. Heatmap shows genes by cluster matrix (arrows represent MDT clusters). **(F)** UMAP representation of all clusters and their identity. **(G-I)** Cellular heterogeneity of the MDT. **(G)** Dendrogram assembled by hierarchical clustering of average variable gene expression for each cluster shows inter-cluster relationships based on transcription. **(H)** MDT body (top) and ring (bottom) clusters. Shown are example marker genes and their UMAP distribution (left) and expression in mouse brain (right). **(I)** Expression distributions of top three marker genes for MDT ring clusters (node 46; clusters 15, 19) and body (node 44; clusters 6, 17, 20). Y-axis depicts log-normalized counts. (**J-L**) Differential expression of voltage gated ion channels in *Necab*-positive ring cells. **(J)** *Necab*-positive cells co-express *Cfos* during tube test (RNAscope FISH). **(K)** *Necab*-positive cells co-express *Cfos* in both rank-1 and rank-3 mice. **(L)** GSEA: effect of social rank on the VGIC gene set. VGICs (black vertical lines) are sorted with all other genes based on differential expression between rank-1 (left) and rank-3 (right). Enrichment score (red line) is a running sum (from left to right) that increases as VGIC genes are encountered and decreases when they are not. Genes left of the yellow line (peak enrichment score) are upregulated in rank-1. The correlation of gene expression with social rank is shown in the bottom histogram (green). Permutation test corrected for multiple testing (FDR), p < 0.01.

We next sought to identify the molecular mechanisms underlying changes in membrane properties of MDT neurons according to social rank. We began by assessing the cellular composition of midline thalamic nuclei using single nucleus RNA-seq (sNuc-Seq), and then investigated transcriptional differences associated with social hierarchy across distinct MDT cell populations (Fig. 4E and SOM Methods). We profiled 33,582 neurons from rank-1, rank-3, and control mice. Sequences of nucleus-barcoded cDNAs were obtained using a droplet microfluidics platform (10x Genomics Chomium), yielding an average of 4,699 transcripts and 2,169 unique genes per nucleus (Fig. 4E). Nuclei were clustered together according to transcriptional similarities using unsupervised principal-component analysis and shared nearest neighbor graph clustering (Macosko et al., 2015; Satija et al., 2015). To assign transcriptional and anatomical identity to the clusters, we identified cluster-specific marker genes based on differential expression between nuclei, and then examined the expression patterns of these markers in the mouse Brain Atlas (Lein et al., 2006). For visualization, the sequence profiles of nuclei were embedded in two dimensions using UMAP (Butler et al., 2018) according to statistically significant PCs and labeled by cluster identity (Fig. 4F).

Our clustering detected well-established and novel thalamic cell populations in which different samples and ranks were homogenously distributed. We identified a specific cluster corresponding to the reticular thalamus, a known inhibitory (GABAergic) population (Krol et al., 2018) that robustly expressed *Gad1*, *Gad2,* and the marker *Isl1* (Fig. 4F and SOM Fig. 4F-G). We also identified five clusters corresponding to the MDT, all of which robustly expressed the excitatory marker *Slc17a6* (VGLUT2) (SOM Fig. 4H). MDT clusters accounted for 12,782 nuclei and were separable into two groups according to transcriptional relationships, UMAP space, and anatomy (Fig. 4G-H). In the cluster dendrogram, three clusters were expressed within the central compartment of the MDT (“body cells”) and associated with expression of the gene *Kcnip3* (Fig. 4H-I). Two clusters were expressed in a ring around the MDT (“ring cells”) and were associated with expression of the gene *Necab1* (Fig. 4 H-I). These results are in agreement with a recent study showing that genes related to potassium channels (i.e. *Kcnip3*) and calcium binding proteins (i.e., *Necab1*) are strongly associated with cell types of the thalamus, including the MDT (Phillips et al., 2019). Thus, our analysis successfully partitioned nuclei of the MDT into putative cell types, providing a point of entry for differential expression analyses.

We next asked how hierarchy formation affected transcription relevant to MDT neuronal structure and function using gene set enrichment analysis GSEA (Mootha et al., 2003, 2004). We tested for differential expression of each gene set or biological pathway (N = 145, see Appendix 1 for gene sets), at each MDT cluster (N = 5), for each of three pairwise comparisons (rank-1 vs. control; rank-3 vs. control; and rank-1 vs. rank-3), for a total of 2,175 tests. Biological pathways that were significant in two or more tests (N=61/2,175) were found in all MDT clusters, and were enriched for genes associated with pre- and post-synaptic structures (46/61, 75%), cellular enzymes including chromatin modifiers and CMGC kinases (16%), voltage gated ion channels (VGICs, 5%), and nuclear hormone receptors (2%) (SOM Fig. 4I).

### Social rank dependent changes to MDT are associated with changes in Trpm3 expression

To further identify genes that were differentially expressed between dominant and subordinate mice, we analyzed the intersection of GSEA results with a genome-wide differential expression test, where each gene passing expression thresholds was tested for an effect of rank for each cluster (SOM Methods). The strongest effects of rank were found in the *Necab1-*positive clusters of the MDT ring (SOM. Fig. 5A). *Necab1* is expressed in a ring encompassing the MDT, PVT, and intralaminar nuclei (Fig. 4J). Moreover, over 80% of *Cfos*+ cells following tube tests were *Necab1*+ (Fig. 4J-K), suggesting that this population is specifically recruited during the display of social hierarchy. Differentially expressed genes were enriched for voltage gated ion channels (Fig. 4L and SOM Fig. 5B) and tyrosine kinase genes (SOM Fig. 5C-D). We identified five voltage gated ion channels that were more highly expressed in rank1 males (*Mcoln2*, *Trpm3*, *Kcnh8*, *Scn3a*, *Cacna1b,* SOM Fig. 5A), and focused further analyses on these genes due to the unequivocal role of ion channels in membrane excitability.

**Figure 5.**
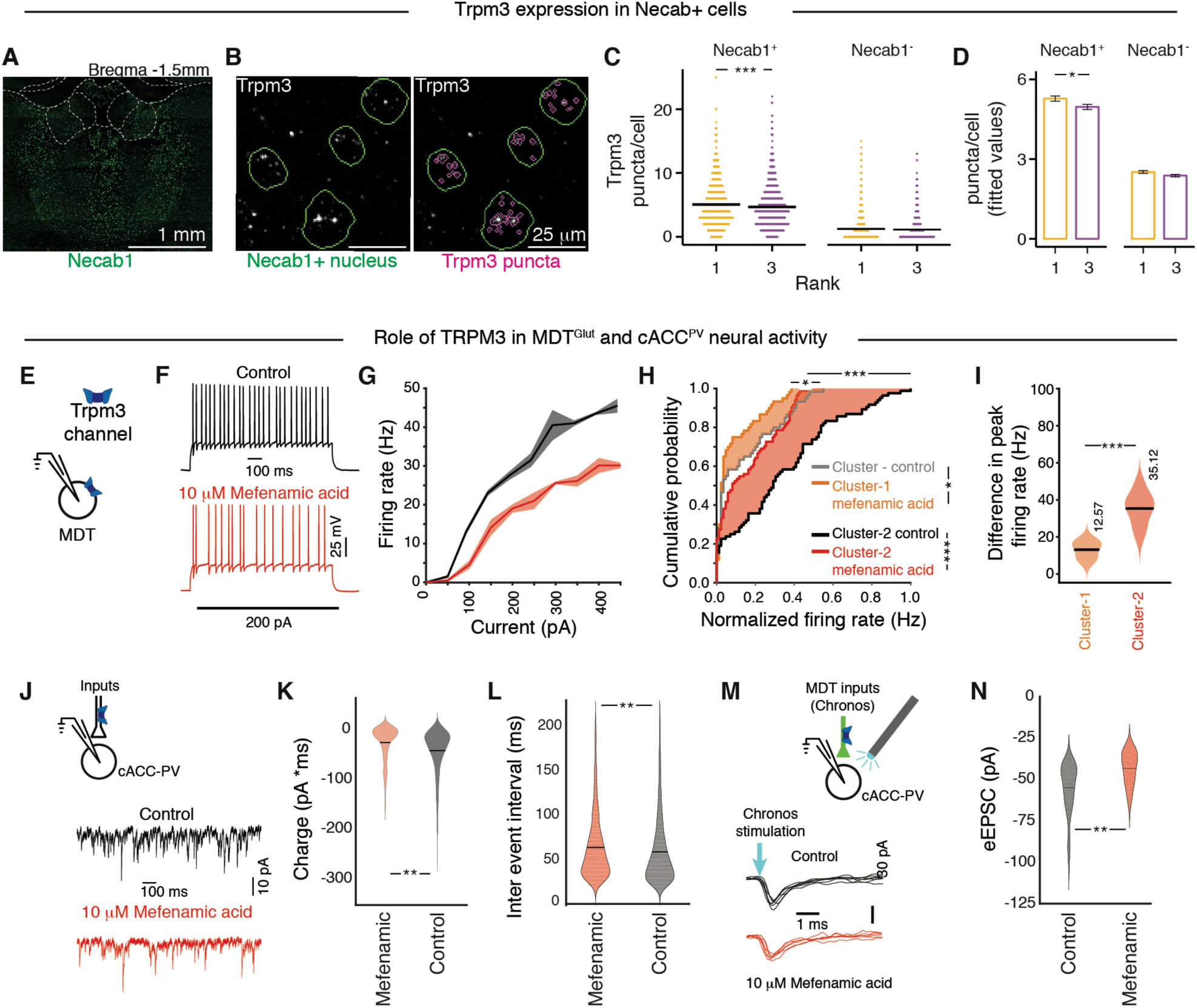
TRPM3 expression in ring cells of the MDT is rank-dependent and enhances neuronal excitability. **(A-D)** RNAscope FISH validation of *Trpm3* expression in *Necab1* ring cells. **(A)** Spatial distribution of *Necab1* cells. **(B)** *Necab1*+ nuclei (green outlines) and *Trpm3* (white speckles and magenta outlines). **(C)** *Trpm3* transcript (puncta) counts in *Necab1*+ cells (N = 35,865) and *Necab1*-cells (N = 69,530). Each dot corresponds to one cell. Horizontal bars show mean and s.e.m for each group. A random subset (50%) of the raw data is plotted to aid in visualization. Wilcox rank sum. **(D)** Fitted values from LMER. **(E-M)** The role of TRPM3 in the MDT-cACC circuit. **(E)** Schematic depicts current-clamp recordings in the MDT. **(F)** Example of firing patterns of MDT neurons under bath application of vehicle (control, black) or TRPM3 antagonist (mefanamic acid, red). MDT frequency/current curves **(G)** and cumulative probability distribution of firing rates **(H)** under control and antagonist conditions for cluster-1 and cluster-2 neurons in MDT, N = 5 for cluster-1 and N = 6 for cluster-2. (I) Summary data depicting the decrease in peak firing rates of cells from cluster-1 (orange) and cluster-2 (red) **(J)** Schematic of cACC^PV^ voltage-clamp recordings. Example sEPSCs in cACC^PV^ cells, colors as in **(F).** Violin plots of **(K)** charge transfer and **(L)** frequency of cACC^PV^ sEPSCs under control and antagonist conditions. **(M)** Schematic of cACC^PV^ voltage-clamp recordings and optogenetic activation of MDT projections. **(N)** Violin plot shows charge transfer for eEPSCs under control and antagonist conditions (n=6 neurons). Kolmogorov Smirnov test, *p<0.05, **p<0.01, ***p<0.001.

We then sought to validate differential expression of voltage gated ion channels in ring cells using quantitative FISH. We first selected *Trpm3*, a transient receptor potential (TRP) channel activated by changes in membrane voltage, temperature, and endogenous neurosteroids including pregnenolone sulfate (Held et al., 2015). As predicted from RNA-seq data, mRNA expression (i.e., number of single molecule RNA FISH puncta) of *Trpm3* in *Necab1* cells was higher in rank1 compared to rank3 (Fig. 5 A-D). As a control, we measured *Trpm3* expression in *Necab1*-negative cells of the MDT and found no effect of rank (Fig. 5 C-D). Thus, upregulation of the *Trpm3* channel is an attribute of social dominance.

### TRPM3 regulates neural excitability in the MDT

We next investigated the role of TRPM3 in the firing rate of MDT neurons using current-clamp recordings (Fig. 5 E-F). TRPM3 is a non-selective cation channel permeable to Ca^2+^, and, to a lesser extent, K^+^, Na^+^ (Held et al., 2015; Oberwinkler et al., 2005). Compared to the control solution, bath application of 10 μM mefenamic acid, a selective TRPM3 antagonist (Klose et al., 2011) caused a marked reduction in the excitability and firing rate of MDT neurons (Fig. 5F-I, p<0.001 for Kolmogorov Smirnov two sample test). Moreover, based on our findings that high-firing cells were relatively more abundant in the MDT of rank-1 than lower rank males (Fig. 4D) and that *Trpm3* expression is upregulated in rank-1 compared to rank-3 males (Fig 5C-D), we predicted that *Trpm3* function should preferentially contribute to the intrinsic properties of cluster-2 (high-firing) compared to cluster-1 (low firing) neurons. Indeed, blocking *Trpm3* function with mefenamic acid resulted in a significant decrease in the firing rates of both populations of MDT neurons (Fig. 5H, p=0.03 for cluster-1 neurons and p<0.001 for cluster-2 neurons, Kolmogorov Smirnov two sample test), with a reduction in peak firing rate substantially more pronounced for neurons in cluster-2 than cluster-1 (Fig. 5I, p < 0.001 for Wilcoxon rank-sum test).

We then tested the hypothesis that presynaptic TRPM3 channels in axon terminals may affect transmitter release, and therefore excitatory synaptic currents recorded in cACC^PV^ cells (Fig. 5I). Using PV-Cre::flex-tdTomato reporter mice to identify PV interneurons in slice (SOM Fig. 5M), we found that mefenamic acid caused a reduction in the frequency and the charge carried by spontaneous excitatory postsynaptic currents (sEPSCs) in PV interneurons (Fig. 5J-L). We then expressed the optical actuator Chronos (AAV5-Syn-Chronos-GFP) in MDT neurons (SOM. Fig. 5 L-M) in PV-Cre::flex-tdTomato reporter mice. Next, we used optogenetics to test whether excitatory transmission in MDT-cACC^PV^ terminals was also dependent on axonally expressed TRPM3 by recording light-evoked currents (eEPSCs) in tdTomato-positive cells (Fig. 5M). Indeed, application of mefenamic acid caused a decrease in the amplitude of excitatory currents in cACC^PV^ interneurons (Fig. 5N).

We conclude that, in MDT ring cells, expression of TRPM3 is upregulated in dominant compared to subordinate mice, promoting high firing rates in MDT neurons and potentiating excitatory release onto cACC^PV^ interneurons.

### Inputs to MDT undergo synaptic plasticity during hierarchy formation

We next investigated changes in synaptic inputs to MDT neurons according to social rank. We measured spontaneous synaptic events in MDT neurons using voltage-clamp recordings. Spontaneous excitatory and inhibitory post-synaptic currents (sEPSCs and sIPSCs) were measured at −70 mV and 0 mV respectively (Fig. 6 A-B). Relative to individually housed controls, rank-1 MDT neurons received more frequent and larger sEPSCs, whereas rank-2 and rank-3 males were similar to controls (frequencies: Fig. 6 C & E, amplitudes: Fig. 6 A & H). In contrast, the frequency of sIPSCs was higher in rank-2 and rank-3 males compared to controls, while rank-1 males were similar to controls (Fig. 6 B, D & F). There was no effect of social rank on the amplitude of sIPSCs (Fig. 6H). We found that rank-1 males received more excitation and less inhibition than rank-2, rank-3 and control males (Fig. 6G). These results suggested that the strength of synaptic inputs to MDT neurons changes in a rank-dependent manner.

**Figure 6.**
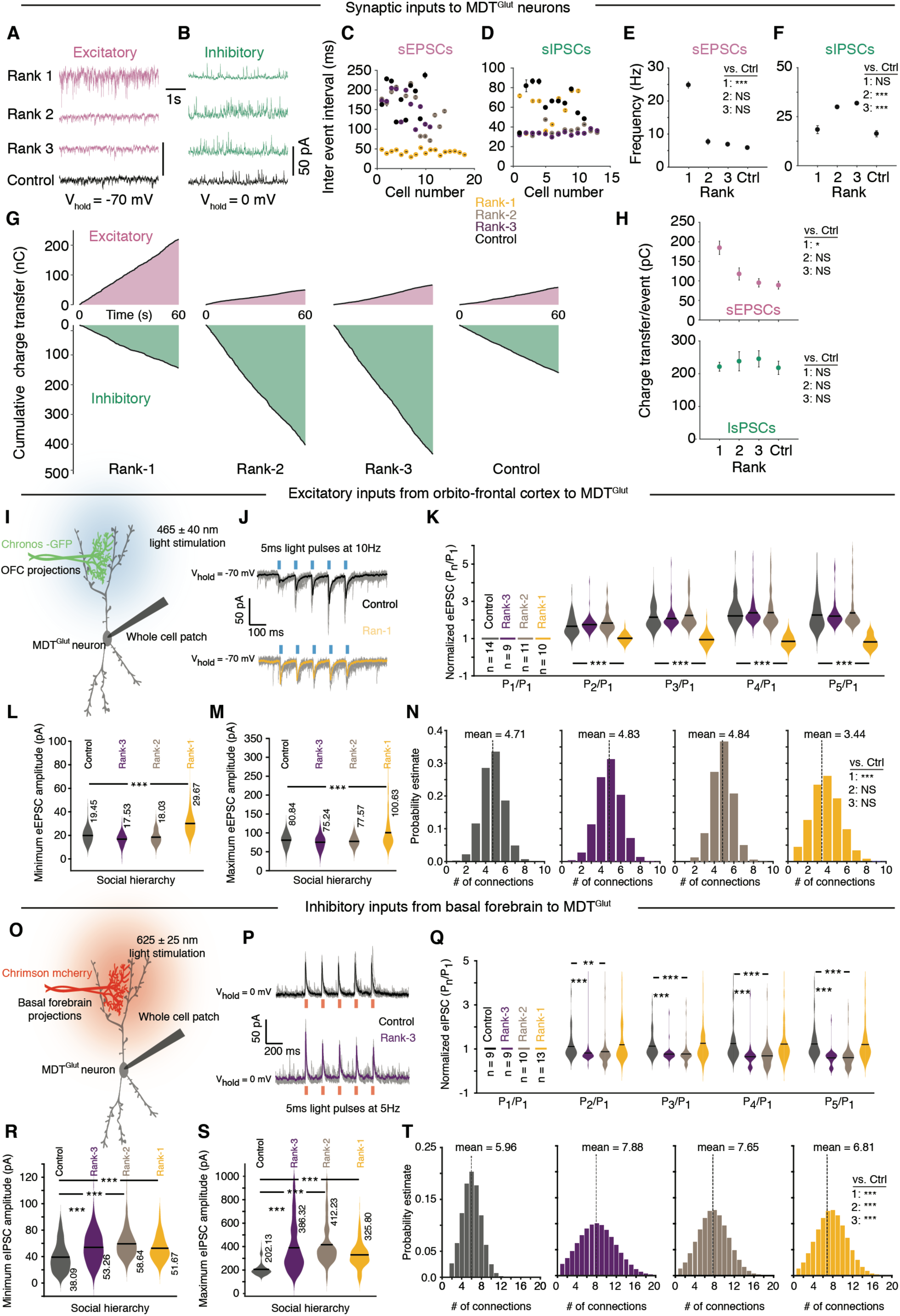
Plasticity of excitatory and inhibitory inputs to medial dorsal thalamus. **(A-F)** Whole-cell voltage-clamp recordings of synaptic activity in MDT^Glut^ neurons. Example sEPSCs **(A)** and sIPSCs (**B**) according to social status. **(C)** Inter-event interval for sEPSCs for all recorded cells by social status. **(D)** Inter-event interval for sIPSCs for all recorded cells by social status. **(E)** Frequency summary of sEPSCs of all cells in **(C).** Inset shows statistical effects relative to controls; means ± s.e.m, N = 53 neurons. **(F)** Frequency summary of sIPSCs of all cells in **(D).** Inset shows statistical effects relative to controls; means ± s.e.m, N = 43 neurons. **(G)** Examples of per-minute cumulative charge transfer of sEPSCs (top) and sIPSCs (bottom) by social status. **(H)** Charge transfer summary: sEPSCs (top) and sIPSCs (bottom), Inset shows statistical effects relative to controls; means ± s.e.m; means ± s.e.m, N = 53 (excitatory event) and 43 (inhibitory event) neurons. **(I)** Schematic depicting electrophysiological recordings of neurons in medial dorsal thalamus (MDT) slices in response to optogenetic activation of orbitofrontal cortex projections (OFC, labelled with AAV chronos). **(J)** Top: electrophysiological traces from control mice (grey: 10 trials, black: mean response of 20 consecutive trials) depicting paired pulse facilitation of excitatory post synaptic activity (eEPSCs) in response to activation of OFC projections (light pulses: 5 ms duration, 465 nm at 10 Hz). Bottom: same paradigm results in paired pulse depression in rank-1 animals. Mean response is shown in yellow. **(K)** Summary of the experiments showing that the control (black), rank-2 (brown) and rank-3 (purple) animals show similar paired pulse facilitation while rank-1 animal show paired pulse depression (N = 9 animals). **(L)** Minimum and **(M)** maximum evoked excitatory responses in MDT neurons in response to activation of OFC projections. **(N)** Histograms showing the distribution of number of excitatory synaptic connections between OFC projections and a single MDT neuron for control (black), rank-1 (yellow), rank-2 (brown) and rank-3 (purple). **(O)** Schematic depicting electrophysiological recordings of neurons in medial dorsal thalamus (MDT) slices in response to optogenetic activation of basal forebrain projections (labelled with AAV chrimson). **(P)** Top: electrophysiological traces (grey: 5 trials, black: mean response of 10 consecutive trials) for paired pulse paradigm for inhibitory post synaptic activity (eIPSCs) in response to activation of basal forebrain projections (light pulses: 5 ms duration, 625 nm at 5 Hz). Bottom: same paradigm results in paired pulse depression in rank3 animals. Mean response is shown in purple. **(Q)** Summary of experiments involving paired pulse paradigm showing that the control (black) and the rank-1 (yellow) animals show neither facilitation nor depression while rank-2 (brown) and rank-3 (purple) animals show paired pulse depression (N = 8 animals). **(R)** Minimum and **(S)** maximum evoked inhibitory responses in MDT neurons in response to activation of basal forebrain projections. **(T)** Histograms showing the distribution of number of inhibitory synaptic connections between basal forebrain projections and a single MDT neuron for control (black), rank-1 (yellow), rank-2 (brown) and rank-3 (purple). ** p<0.01 and *** p<0.001.

To determine the identity of rank-dependent excitatory or inhibitory inputs to the MDT, we next assessed plasticity in functional connectivity at MDT synapses using optogenetics (SOM Fig. 6 A-B). Short-term plasticity refers to the alteration of synaptic strength caused by patterns of presynaptic firing activity on the order of milliseconds and is critical for information processing and computation at synapses (Regehr, 2012). Short-term plasticity is measured as post-synaptic responses to repeated presynaptic stimulation and can be observed as either increases or decreases in synaptic strength following the first stimulus (i.e., facilitation or depression). We injected channelrhodopsin (ChR2) variants Chronos or Chrimson (SOM Fig. 6B) in four areas presynaptic to the MDT (piriform cortex, lateral preoptic area, OFC, and basal forebrain) that met two criteria: (1) identified in viral tracing experiments (Fig. 3 A-C), and (2) positive for CTB-retrograde labeling and CFOS immunohistochemistry following the tube test (data not shown). We then recorded optically evoked postsynaptic currents (eEPSCs and eIPSCs) in MDT neurons using 5 ms pulses (5 or 10 Hz) (Fig. 6I-J and SOM Methods).

We used a paired pulse stimulation protocol to estimate the probability of release from inputs to the MDT, and to compare data from animals of different ranks (Fig. 6J) (Jackman et al., 2016). We found that the OFC made excitatory synapses with MDT neurons (Fig. 6J). Control and lower rank (rank-2 and rank-3) animals showed significant facilitation (p < 0.01 for Wilcoxon rank sum test, Fig. 6J-K) during paired pulse stimulation paradigm (SOM Methods), suggesting that these synapses have low probability of release. By contrast, we found that the same synapses in dominant animals showed significant depression during paired pulse stimulation (p< 0.01 for Wilcoxon rank sum test, Fig. 6J-K), suggesting a higher probability of release from OFC projections in rank-1 animals. To further document the potentiation of OFC to MDT synapses in dominant animals, we measured the minimum and the maximum amplitudes of OFC eEPSCs (SOM Methods). Dominant animals showed a significant increase in the minimum eEPSCs (Fig. 6L, control = 19.47 pA, rank-3 = 17.53 pA, rank-2 = 18.03 pA and rank-1 = 29.67 pA, p < 0.001 for Wilcoxon rank sum test) and maximum eEPSCs (Fig. 6M, control = 80.84 pA, rank-3 = 75.24 pA, rank-2 = 77.57 pA and rank-1 = 100.63 pA, p< 0.001 for Wilcoxon rank sum test). These results suggest that OFC inputs to MDT neurons are preferentially potentiated in dominant animals and that this is achieved by both increasing the probability of release (Fig. 6J-K) and size of the minimum release event (Fig. 6L).

Next, we estimated the number of inputs by comparing minimum and maximum amplitudes of eEPSCs (Litvina and Chen, 2017). We found that rank-1 animals had significantly fewer connections between OFC and MDT (Fig. 6N, 3.44 connections for rank-1 and 4.71, 4,83 and 4.84 connections for control, rank-3 and rank-2 animals respectively, p<0.001 for Wilcoxon rank sum test). In dominant animals the synapses between the OFC and MDT neurons undergo significant change, resulting in fewer (Fig. 6N), but stronger, individual synapses than those in the lower ranks (Fig. 6L-N).

Using a similar optogenetic strategy (Fig. 6O and SOM Fig. 6D), we tested the inputs from basal forebrain (nucleus of the diagonal band (NDB) and substantia innominata) to MDT. By recording eEPSCs at −70 mV (SOM Fig. 6 D) and eIPSCs at 0 mV (Fig. 6P), we found that the basal forebrain sends both excitatory and inhibitory inputs to MDT neurons. Intriguingly, inhibitory and excitatory projections showed remarkably distinct synaptic properties. Inhibitory projection showed neither facilitation nor depression in control animals (Fig. 6P-Q), whereas lower ranks (rank-2 and rank-3) showed significant paired pulse depression during paired pulse paradigm (p<0.001 for Wilcoxon rank sum test, Fig. 6Q, rank-1 was not significantly different from control). Since increased paired pulse depression is typically associated with increased release probability (Regehr, 2012), these data show that the inhibitory inputs from basal forebrain to MDT are significantly potentiated in rank-2 and rank-3 animals. In contrast, excitatory projections showed paired pulse depression (SOM Fig. 6D-E), and the extent of depression was unaffected by the social status of the animal (SOM Fig. 6D-E). Like our previous experiment, we measured the minimum and maximum amplitude of eIPSCs and eEPSCs in MDT neurons in response to optogenetic stimulation of the basal forebrain projections. We observed that irrespective of rank, inhibitory inputs were significantly potentiated (Fig. 6R-S, for minimum IPSC amplitude, control = 38.09 pA, rank-3 = 53.26 pA, rank-2 = 58.64 pA and rank-1 = 51.67 pA and for maximum eIPSC amplitude, control = 202.13 pA, rank-3 = 386.32 pA, rank-2 = 412.23 pA and rank-1 = 325.80 pA). This potentiation was higher for the rank-2 and rank-3 as compared to rank-1 animals. In contrast, excitatory inputs from basal forebrain to MDT showed no such change (SOM Fig. 6F-G, for minimum eEPSC amplitude, control = 21.76 pA, rank-3 = 20.15 pA, rank-2 = 18.88 pA and rank-1 = 20.63 pA and for maximum eEPSC amplitude, control = 149.43 pA, rank-3 = 150.75 pA, rank-2 = 151.48 pA and rank-1 = 155.12 pA). These results suggest that the BF inputs to MDT neurons are preferentially potentiated in subordinate animals and that this is achieved by both increasing the probability of release (Fig. 6P-Q) and size of the minimum release event (Fig. 6L).

Using the method described above, we quantified the number of connections between the basal forebrain and MDT neurons for both inhibitory and excitatory inputs. The number of inhibitory synapses increased significantly in animals that have undergone hierarchy training compared to control animals (Fig. 6T, p<0.001 for all ranks, Wilcoxon rank sum test). This increase was higher for the rank-2 and rank-3 animals compared to rank-1 animals. Taken together, these results (Fig. 6G & 6I-T) suggest that the acquisition of social rank through our three step paradigm (Fig. 1G) results in remarkable synaptic plasticity (Fig. 6L-N & 6R-S), with increased excitatory drive and decreased inhibitory inputs in the MDT of dominant animals. In contrast, the MDT of subordinate animals receives higher inhibitory inputs but lower excitatory inputs.

Thus, persistent changes to the intrinsic properties of MDT neurons, and the strength of connections at MDT synapses, occurs in both high rank and low rank males during hierarchy establishment. Together with our *Cfos* data (Fig. 2A) and chemogenetic data (Fig. 2L), these results suggest that electrophysiological properties of MDT neurons are shaped during the formation of social hierarchy to establish circuit-level differences associated with social rank.

## DISCUSSION

We have used behavioral analysis along with a suite of molecular and genetic tools to further our understanding of how a functional neural circuit is altered as social hierarchy emerges. This work leads to several significant advances. (1) We developed a paradigm to quantify hierarchy formation among genetically similar mice and identified key behavioral and physiological attributes of social rank acquisition. (2) We identified interconnected brain areas that are differentially activated in dominant and subordinate mice, including the MDT-ACC circuit. (3) We found that MDT is essential for hierarchy formation while cACC is crucial for retention and maintenance of established hierarchy(4). We uncovered how intrinsic and synaptic properties of MDT neurons, including the strength of inputs from presynaptic OFC and BF neurons to MDT and the strength of MDT inputs to cACC^PV^ neurons, changes as animals become dominant or subordinate. (5) We identified transcriptional changes in MDT ring cells that are associated with the emergence of hierarchy, among which an increase in expression of the voltage-gated ion channel *Trpm3* in dominant males, which enhances MDT neuronal excitability and potentiates synaptic inputs to cACC^PV^ interneurons. Altogether, these findings significantly enhance our understanding of the neural mechanisms underlying the acquisition and maintenance of social rank.

### A model for the functional role of MDT in social hierarchy behavior

The MDT has been proposed to regulate social hierarchy through its connectivity with dmPFC based on the findings that winning could be triggered by excitation of dmPFC pyramidal cells, or by photostimulation of MDT terminals in dmPFC (Wang et al., 2011; Zhou et al., 2017). These observations suggested a feedforward excitation model, where MDT excitation causes excitation of dmPFC pyramidal cells and their downstream targets such as the Nucleus Accumbens, thus leading to increased competitive motivation and reward in high rank males. By contrast, our results point to a separate, though not mutually exclusive, cortical feedforward inhibition mechanism where, in high rank males, MDT^Glut^ neurons, through their projection to cACC^PV^ inhibitory interneurons, may decrease activity of cACC pyramidal cells and thus of downstream circuits such as the amygdala-dependent fear pathways (Toyoda et al., 2011). Consistent with this model, the brain wide *Cfos* mapping done in our study showed opposing patterns of neuronal activity in MDT (up in dominants) and cACC (up in subordinates), while our correlational analysis uncovered the MDT and cACC as two particularly divergent nodes in a brain-wide network. We showed that MDT^Glut^ projections synapse onto cACC^PV^ interneurons and that MDT-cACC projections are differentially activated between dominants and subordinates. Our chemogenetic experiments demonstrated that competitive performance in tube tests can be enhanced by activation of MDT, or by inhibition of the cACC via excitation of PV interneurons.

Converging evidence from different studies shows that MDT activates inhibitory interneurons of the frontal cortex (Ferguson and Gao, 2018; Miller et al., 2017), including the ACC (Delevich et al., 2015), and that functional roles of this circuity include the control of reward-seeking strategies (Kvitsiani et al., 2013; Sparta et al., 2014) and fear expression (Courtin et al., 2013). Intriguingly, increased firing of ACC^PV^ cells encodes longer wait times during a foraging task (Kvitsiani et al., 2013) and reduces fearful behavior in a fear conditioning task (Courtin et al., 2013). In addition, a previous study found that mice subjected to repeated social defeat exhibited heightened social avoidance and weakened functional connectivity in the MDT-mPFC circuit (Franklin et al., 2017). In light of these studies, our results suggest a scenario in which MDT activity may instruct the cACC to reduce fear expression during head-to-head encounters. Indeed, activation of the MDT increased competitive performance of subordinate males, but also resulted in less anxious behavior in the open field test. Thus, a more general interpretation of the role of the MDT may be to engage attentional processing by reducing the expression of anxiety-related behaviors.

Our results also revealed an intriguing interplay between MDT and cACC in shaping and maintaining social structure. We found that MDT function is essential for the establishment of social hierarchy but dispensable after animals had learned their rank, while cACC plays a critical role in the retention of the established hierarchy. In established hierarchies, suppressing activity of cACC pyramidal cells by exciting PV interneurons resulted in increase in the social status of subordinate animals. In contrast, disinhibiting cACC pyramidal cells by inhibiting PV interneurons resulted in drastic loss of social standing of the dominant animals. Intriguingly, in established hierarchies, while cACC was important to preserve the memory of individual’s social position, MDT still retained the ability to maintain influence over individuals’ social rank. In subordinate animals in well-established social groups, we were able to selectively chemogenetically stimulate the activity of MDT neurons, thus suppressing the activity of cACC pyramidal cells. This manipulation led to substantial increase in the social rank of the subordinate individuals. In contrast, MDT lesions in established hierarchies had no effect on the social ranks.

Taken together, these results suggest that the loss of function in MDT once downstream circuits have learned has no effect on social hierarchy, while gain of function—activation—of MDT even in already established social groups has profound effects. By contrast, bidirectional manipulation of cACC activity has an effect on pre-established social hierarchy. These data suggest that memory of rank is stored downstream of MDT, which may at least partially occur in the cACC.

### Linking social status with MDT cell-type specific transcription and circuit physiology

Although the morphological and hormonal basis of social rank is increasingly well understood (Sapolsky, 2004), a long-standing question remains as to how neural circuits evolve at the molecular and cellular level as social rank emerges (Qu et al., 2017; Wang et al., 2014). Here we provide mechanistic insights at the biophysical, molecular, and cell type-specific levels on how circuits change according to social rank.

Transcriptome analysis of over 12,000 single nuclei from the MDT delineated two major cell populations: ring cells and body cells. While exploring rank-dependent transcriptional differences, we found a set of candidate voltage gated ion channels that were differentially expressed between dominants and subordinates, a finding concordant with our observation that MDT neurons were more excitable in dominant males. In particular, the calcium-permeable cation channel *Trpm3* was more highly expressed in MDT ring cells of dominant males. Using a combined electrophysiology and pharmacology approach, we found that blocking TRPM3 channels decreased MDT neuronal membrane excitability and excitatory neurotransmission onto cACC^PV^ interneurons. These results suggest a mechanism in which the increase in *Trpm3* expression in dominant males leads to a persistent increase in MDT-ACC feedforward inhibition with significant behavioral consequences. *Trpm3* can be activated by a number of stimuli, including voltage (Held et al., 2018) and pregnenolone sulfate (PregS), a neurosteroid that has been linked to aggression (Baulieu, 1998; Held et al., 2018). Future studies should shed light on how TRPM3 expression is regulated during social competition.

### Emergence of dominant and subordinate status is associated with synaptic plasticity

In addition to changes in the intrinsic excitability of MDT neurons, our experiments revealed remarkable changes in the synaptic connectivity of MDT neurons during hierarchy formation. We identified two brain areas, the orbitofrontal cortex and the basal forebrain, as major sources of excitatory and inhibitory inputs to MDT, respectively, that undergo significant plasticity during acquisition of social rank (Fig. 7). Emergence of dominant behavior is accompanied by strengthening of excitatory inputs from OFC to MDT neurons while emergence of subordinate status is accompanied by strengthening of inhibitory inputs from BF. Our results also revealed significant changes to the membrane properties of MDT neurons, resulting in higher excitability of MDT neurons during attainment of dominant rank. Together, these results show that in dominant animals, MDT receives higher excitatory drive and is also primed to respond more strongly to excitatory inputs. In contrast, subordinate animals receive higher inhibitory inputs and respond comparatively weakly to excitatory inputs. Thus, acquisition of dominant status is accompanied by enhanced MDT-ACC feedforward inhibition while acquisition of subordinate status is accompanied by disinhibition of ACC, which may lead to the enhancement of fearful behavior (Fig. 7). Whether TRPM3 mediated excitability of MDT neurons and presynaptic plasticity are mechanistically linked is not clear. One intriguing possibility is that the level of activity of MDT neurons regulates the strength of both excitatory and inhibitory inputs, as has been demonstrated in cortical pyramidal neurons (Xue et al., 2014).

**Figure 7.**
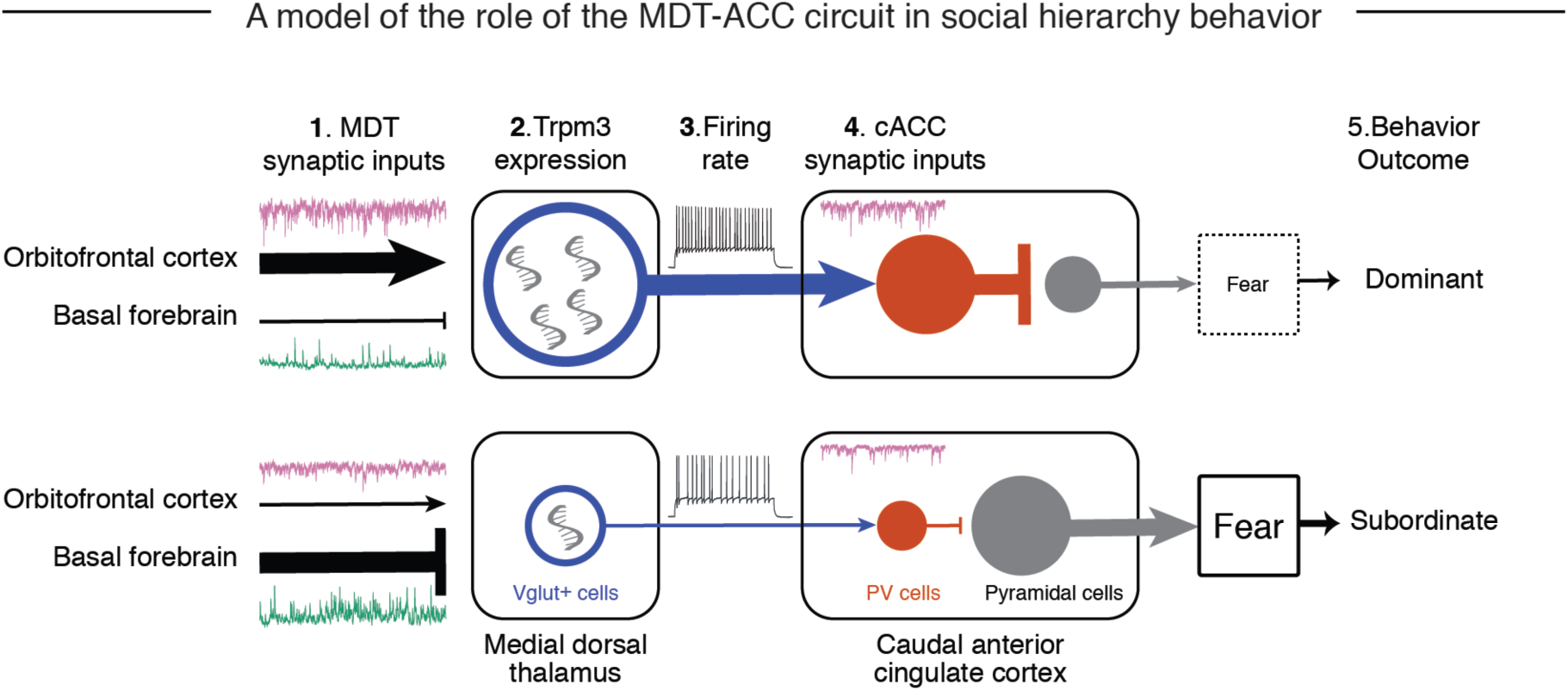
Model: functional role of MDT-cACC circuit in social hierarchy behavior. Compared to subordinates, MDT neurons in dominant males show (1) enhanced spontaneous excitatory activity (sEPSCs, the orbital frontal cortex is the major source of these inputs), and reduced inhibitory activity (sIPSCs, basal forebrain is one source of these inputs), (2) greater expression of the Ca2+-permeable nonselective cation channel Trpm3, and (3) higher firing rates. MDT neurons send excitatory monosynaptic projections to Parvalbumin-positive inhibitory neurons in caudal part of anterior cingulate cortex, and these inputs are (4) enhanced by Trpm3. The enhanced MDT-mediated feedforward inhibition of the cACC in high rank animals drives social dominance behavior perhaps in part through reduction of activity in downstream amygdala fear circuits (5). By contrast, MDT neurons in subordinate males show (1) enhanced spontaneous inhibitory activity (sIPSCs from basal forebrain) and reduced excitatory activity (sEPSCs, from OFC) (2) lower expression of the Ca2+-permeable nonselective cation channel Trpm3, and (3) lower firing rates. This reduced activity of MDT neurons in low rank males leads to disinhibition of cACC pyramidal cells, which may increase activation of downstream fear circuits and drives social subordinate behavior (5)

In summary, our results show that the membrane and synaptic properties of MDT neurons are modulated during hierarchy formation to establish circuit-level differences associated with social rank. These data provide new mechanistic insight into the molecular, physiological and circuit basis of social hierarchy and show how defined changes in excitability at a specific corticothalamic loop may affect complex social interactions. This work opens new avenues to understand how cortical circuits are remodeled to control state-dependent social behaviors.

## Acknowledgements

We thank members of the Dulac and Murthy laboratories for thoughtful discussions and helpful comments on the manuscript, Doug Richardson, Harvard Center for Biological Imaging, Lai Ding, and Jeff Farrell for help with microscopy and image analysis, Stacey Sullivan and the Harvard BRI for help with animal husbandry, Christof Neumann for assistance with the Elo Rating analysis. Yasin Kaymaz and Harvard FAS Informatics for assistance with gene set enrichment analysis. This work was supported by HHMI (CD) and NIH grants R01MH113094, R01MH111502 and U19MH114821 (CD) and R01 DC014453 (VM).

## SUPPLEMENT FIGURE LEGENDS

**SOM Figure 1.**
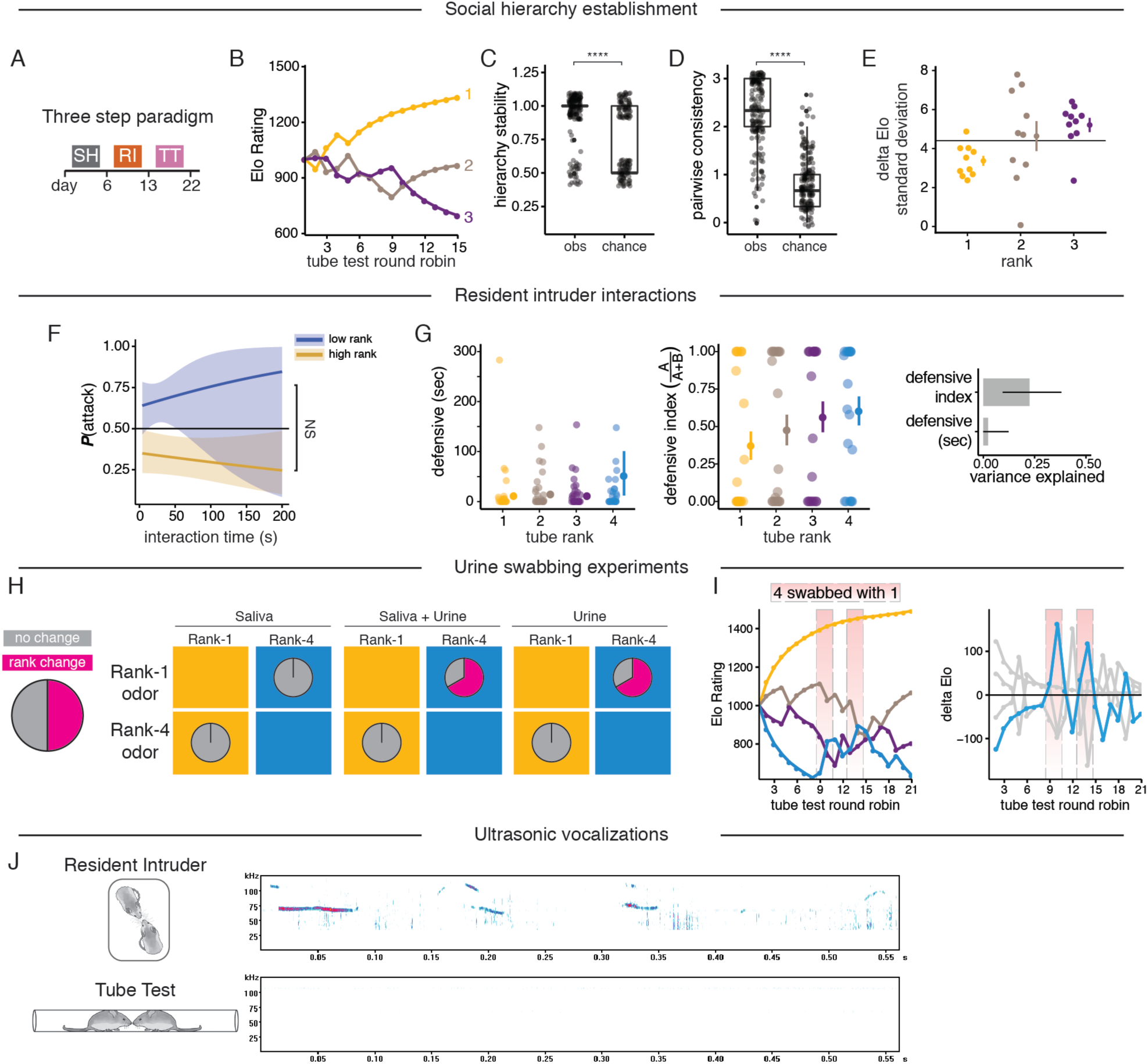
Determinants of hierarchy formation among groups of unfamiliar mice. **(A-E)** Social hierarchy formation within groups of three males. (A) Paradigm for the emergence of social hierarchy among unfamiliar mice. SH: single housed; TT: tube test; RI: resident-intruder. **(B)** Elo ratings and ordinal ranks of an example group of 3 mice tested across 15 round robins. Tube test hierarchy stability **(C)** and pairwise consistency **(D)** of observed groups compared to random outcomes, box plots show median and first and third quartiles. **(E)** Standard deviation in delta Elo scores by tube test rank. **(F)** Probability of attacking behavior as a function of resident-intruder interaction time for high rank and low rank mice. Binomial logistic regression ± s.e.m. **(G)** Time spent displaying defensive behavior (left) and defensive index (middle) as predictors of tube rank; mean ± s.e.m. Right: R-squared ± 95% c.i. represents variance in tube rank explained by defensive index and duration. (G) Urine swabbing experiment transferring saliva, urine or both from rank-1 to rank-4 and vice versa. Rank-4 males gained higher rank when swabbed with rank1 urine. N = 4 social groups. **(H)** Example Elo (left) and delta Elo (right) from a group of four when rank-4 was swabbed with urine from rank-1 (shaded rectangles). **(I)** Ultrasonic vocalizations emitted during resident-intruder (bottom) but not tube test (top). N = 2 groups (8 mice) tested.

**SOM Figure 2.**
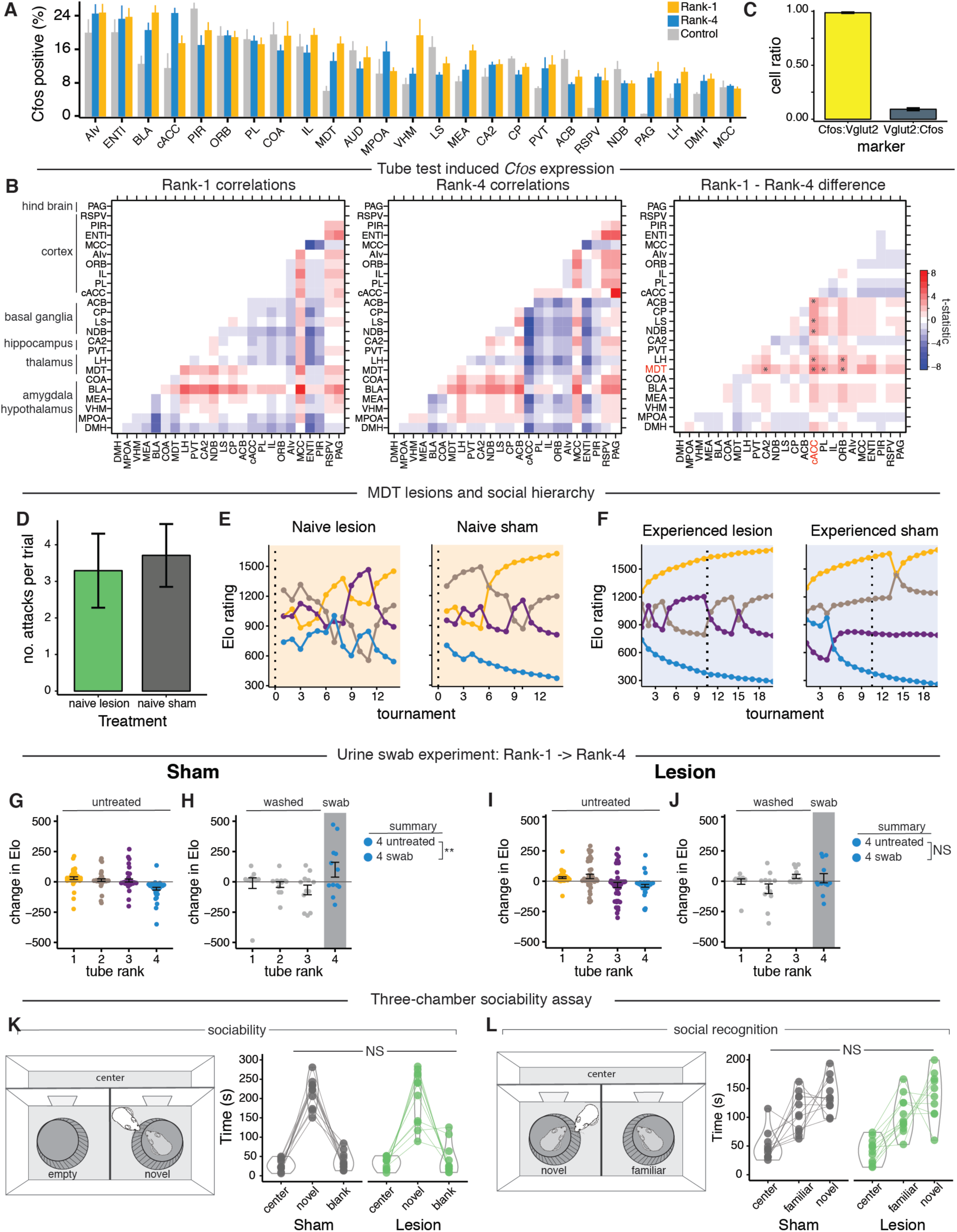
Social rank-dependent brain activity and requirement of MDT in emergence of social hierarchy. **(A)** Quantification of brain-wide *Cfos* expression in rank-1, rank-4, and control males. Mean ± s.e.m. LMER with Tukey post hoc test. N = 4 social groups plus two control males (10 mice). Equivalent to Fig. 2A. **(B)** Correlation matrices of *Cfos* expression across brain regions in rank-1 and rank-4 mice (left and middle), and their difference (right), N = 4 social groups. **(C)** Ratio of *Cfos*-positive to *Vglut2*-positive neurons (yellow) and ratio of *Vglut2*-positive to *Cfos*-positive neurons (gray) in the MDT following tube test interactions. N = 11,849 *Vglut2-*positive cells (2 mice). **(D)** Frequency of aggressive behavior during resident-intruder interactions among the naïve cohorts. **(E-F)** Effects of MDT lesion on hierarchy dynamics. **(E)** Elo rating of naïve animals in tube test (left) MDT lesion and sham control (right). **(F)** Elo rating of experienced animals in tube test (left) MDT lesion and sham control (right). **(G-J)** Effect of MDT lesion on pheromonal signaling and performance (change in Elo). Baseline performance by rank in untreated (non-swabbed) sham **(G)** and lesion **(I)** groups. In swab experiments, all mice were washed, and rank-4 swabbed with rank-1 urine; sham groups **(H)** and lesion **(J)** groups, LMER with Tukey post hoc test, mean ± s.e.m, N = 6 social groups (24 mice), **p<0.01, NS = not significant. **(K-L)** Effect of MDT lesion vs. sham controls on sociability **(K)** and social memory **(L)** on time spent in the three-chamber assay. Spaghetti plots connect same individuals across rooms of the three-chamber apparatus; violins show density and distribution of data, LMER, N = 6 social groups (24 mice). NS = not significant.

**SOM Figure 3.**
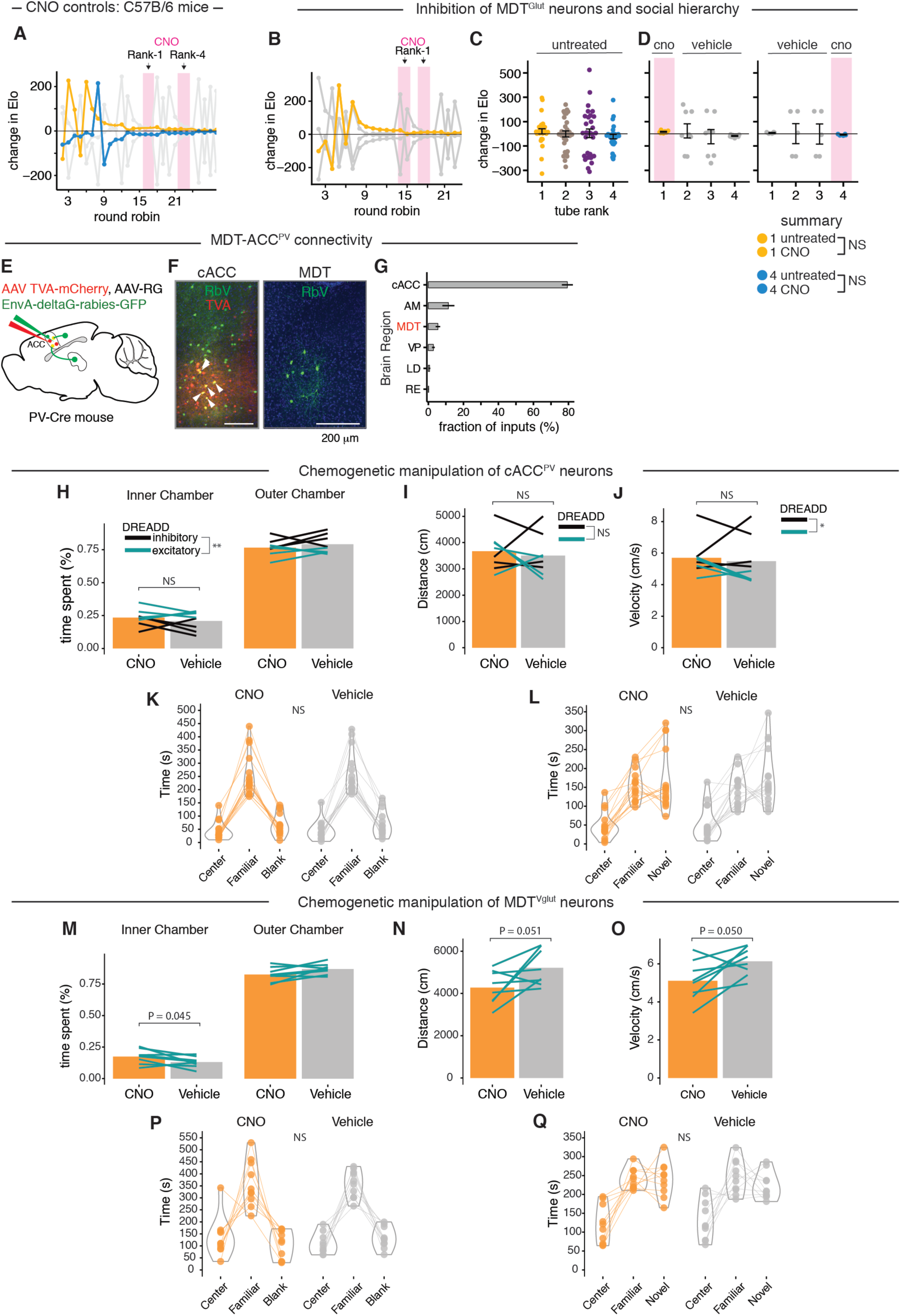
Functional role of cACC^-PV^ and MDT^-Glut^ neurons. **(A)** Effect of CNO alone on social rank in control animals. **(B-D)** Effect of inhibition of MDT^Glut^ on social rank. **(B)** Individual performance (change in Elo) in a hierarchy where the rank-1 male (yellow trace) received CNO (shaded rectangles). Gray lines represent the other three males. **(C)** Relationship between performance and final tube rank in untreated mice. **(D)** Effect of CNO in experiments where either rank-1 (left) or rank-4 (right) received CNO while the other three mice received vehicle. **(E-G)** Monosynaptic retrograde tracing from cACC^PV^ neurons. **(E)** Tracing scheme. **(F)** Overview of cACC (left) and labeling of rabies virus (green), helper AAVs (red), and starter neurons (white arrows) (left). Rabies-positive input neurons in the MDT (right). **(G)** Input fractions to cACC^PV^ neurons (percent of total counted inputs), mean ± s.e.m, N = 3 mice. **(H-J)** Effects of chemogenetic manipulation of cACC-PV neurons in the open field test, (N = 8 mice, means ± s.e.m.), **p<0.01, NS = not significant. **(H)** time spent in open field. **(I)** Distance travelled. **(J)** Velocity. Also shown are effects of the DREADD expressed (inhibitory: black; excitatory: blue-green). **(K-L)** Effect of chemogenetic manipulation on social behaviors, N = 16 mice, means ± s.e.m. **(K)** Sociability assay. **(L)** Declarative social memory assay. **(M-O)** Effects of chemogenetic excitation of MDT^Glut^ neurons. N = 8 mice, mean ± s.e.m. Panels are same as for panels I-M. Statistical effects are from LMER models.

**SOM Figure 4.**
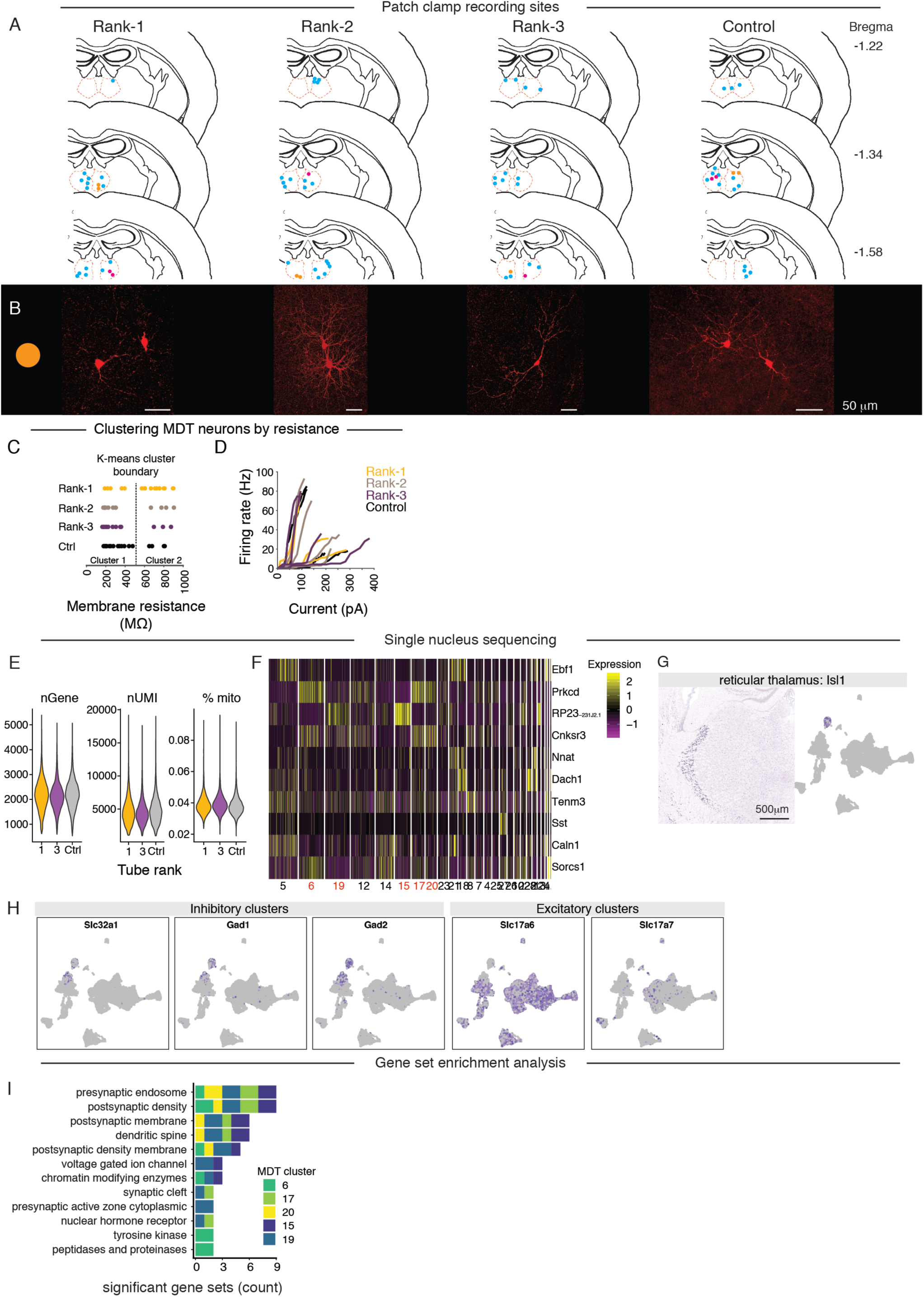
Intrinsic properties of MDT neurons. **(A)** Location of recorded cells. Each dot represents a biocytin filled cell that was recorded, according to social rank and bregma coordinate. In total, 60 neurons were recorded in current clamp, and 53 neurons were recorded in voltage clamp. **(B)** Confocal images of representative biocytin-filled neurons colored according to anatomical coordinates (magenta and gold circles) and labeled by mouse identity. **(C)** K-means clustering of recorded cells based on membrane resistance and according to social rank. Cluster-1: low resistance; Cluster-2: high resistance. **(D)** FI (frequency/current) curves for selected cluster-1 and cluster-2 neurons shown in (C); two neurons per rank per cluster-type were selected. **(E)** Quality control. Shown are the per-nucleus number of genes, UMIs, and percent mitochondria for rank-1, rank-3, and control mice. **(F)** Single nucleus heatmap of marker expression across thalamic nuclei **(G)** The gene *Isl1* corresponds to the reticular thalamus in the mouse brain (left) and is a unique marker for cluster-27 (right). (H) UMAP position of genes encoding markers of inhibitory neurons (*Slc32a1*, *Gad1*, *Gad2*) and excitatory neurons (*Slc17a6*, *Slc17a7*). **(I-J)** Effects of social rank on gene set enrichment in MDT clusters. Gene sets (N = 145) are a curated list representing neural physiology, structure and function. N = 12,782 cells. **(I)** Gene sets affected by social status. X-axis: significant gene set count, where each count is one gene set per cluster per comparison (rank-1 vs. control; rank-3 vs. control, rank-1 vs. rank-3). Colors depict MDT cluster identity.

**SOM Figure 5.**
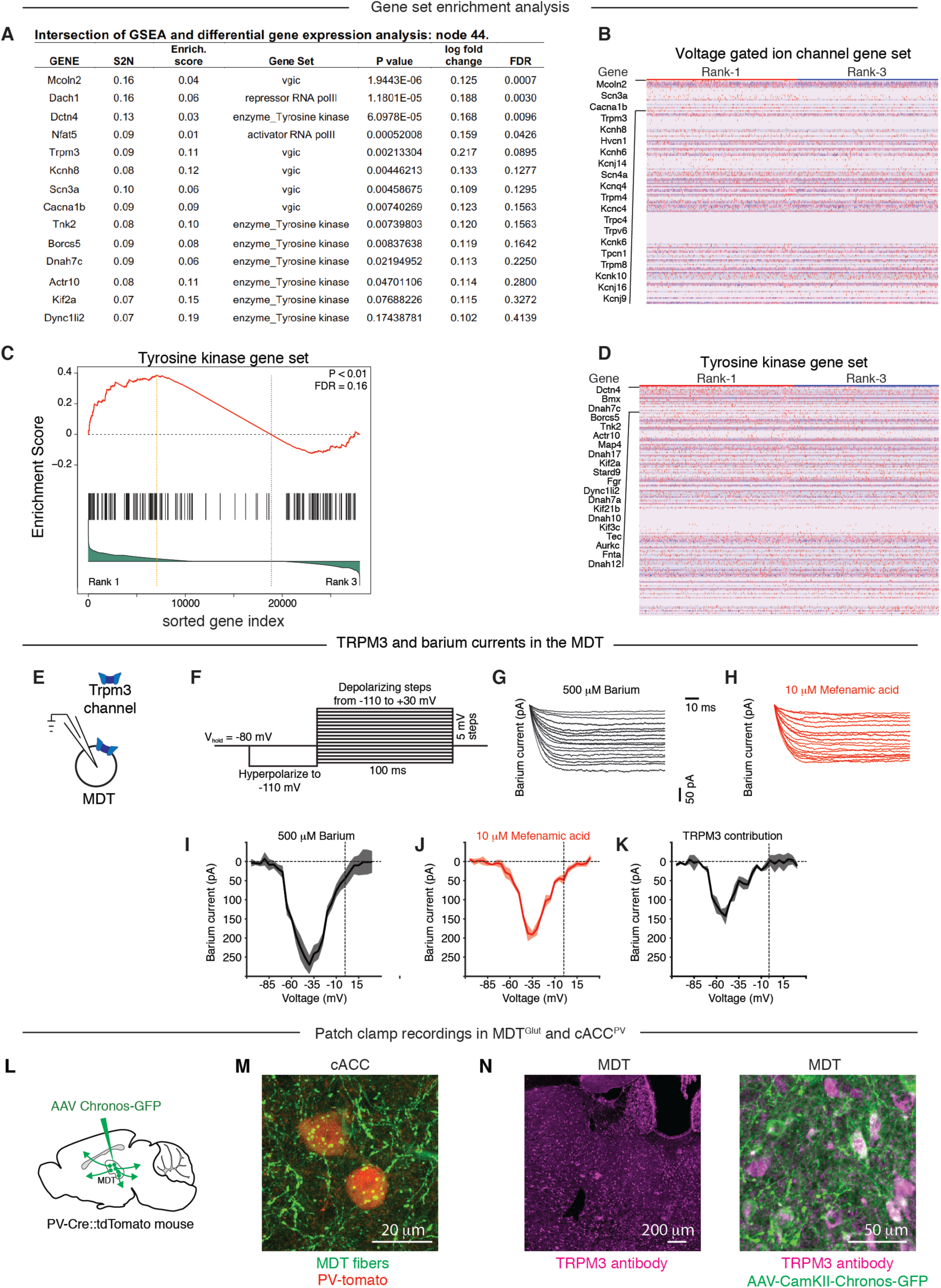
TRPM3 expression in ring cells of the MDT. **(A)** Candidate genes differentially expressed between dominant and subordinate mice in MDT ring cluster cells. Table shows genes at the intersection of gene set enrichment analysis (GSEA) and differential gene expression analysis. S2N: signal to noise ratio, uses the difference of mean expression level (between dominant and subordinate), scaled by the standard deviation. The larger the signal-to-noise ratio, the larger the differences of the means. **(B)** Heatmap showing expression levels of all genes in the voltage-gated ion channel (VGIC) gene set (rows), ordered by the most differentially expressed (top). Each column is a single nucleus. The range of colors (red, pink, light blue, dark blue) shows the range of differential expression values (high, moderate, low, lowest). **(C)** GSEA plot showing effect of social rank on the tyrosine kinase gene set. Tyrosine kinases (black vertical lines) are sorted with all other genes based on differential expression between rank-1 (left) and rank-3 (right). The enrichment score (red line) is a running sum statistic that increases as tyrosine kinase genes are encountered and decreases when they are not, progressing from the strongest to weakest differentially expressed gene. Genes to the left of the yellow line (peak enrichment score) are upregulated in rank-1. The correlation of gene expression with social rank is shown in the bottom histogram (green). Permutation test corrected for multiple testing (FDR), P < 0.01. **(D)** Heatmap showing expression levels of all genes in the tyrosine kinase gene set, as in (B). **(E-K)** TRPM3 is permeable to barium and calcium in the MDT. **(E)** Schematic depicts the slice recording setup. **(F)** Voltage-clamp recording protocol. Barium currents were recorded using a series of depolarizing test pulses (100 ms duration, from −100 to +30 mV, 5 mV increments). Typical electrophysiological traces of inward barium currents in the presence of sodium and potassium blockers **(G)**, plus the TRPM3 antagonist mefanamic acid **(H)**. Note that, unlike calcium currents, barium currents do not inactivate during the long test steps. **(I-K)** TRPM3 contribution to the voltage-current relationship of MDT neurons. Mean voltage-current values of MDT neurons the presence of sodium and potassium blockers **(I)**, plus the TRPM3 antagonist mefanamic acid **(J). (K)** Current subtraction of (I) and (J) reveals TRPM3 contribution, N=5 neurons. **(L)** Schematic depicting injection of an optical actuator Chronos (AAV5-Syn-Chronos-GFP) in MDT neurons in PV-Cre::flex-tdTomato reporter mice. **(M)** MDT fibers labelled with Chronos (AAV5-Syn-Chronos-GFP) in cACC, cACC^PV^ neurons are shown in red**. (N)** Trpm3 antibody staining in MDT^glut^ neurons (left) and overlap of Chronos-GFP and Trpm3 antibody staining (right).

**SOM Figure 6.**
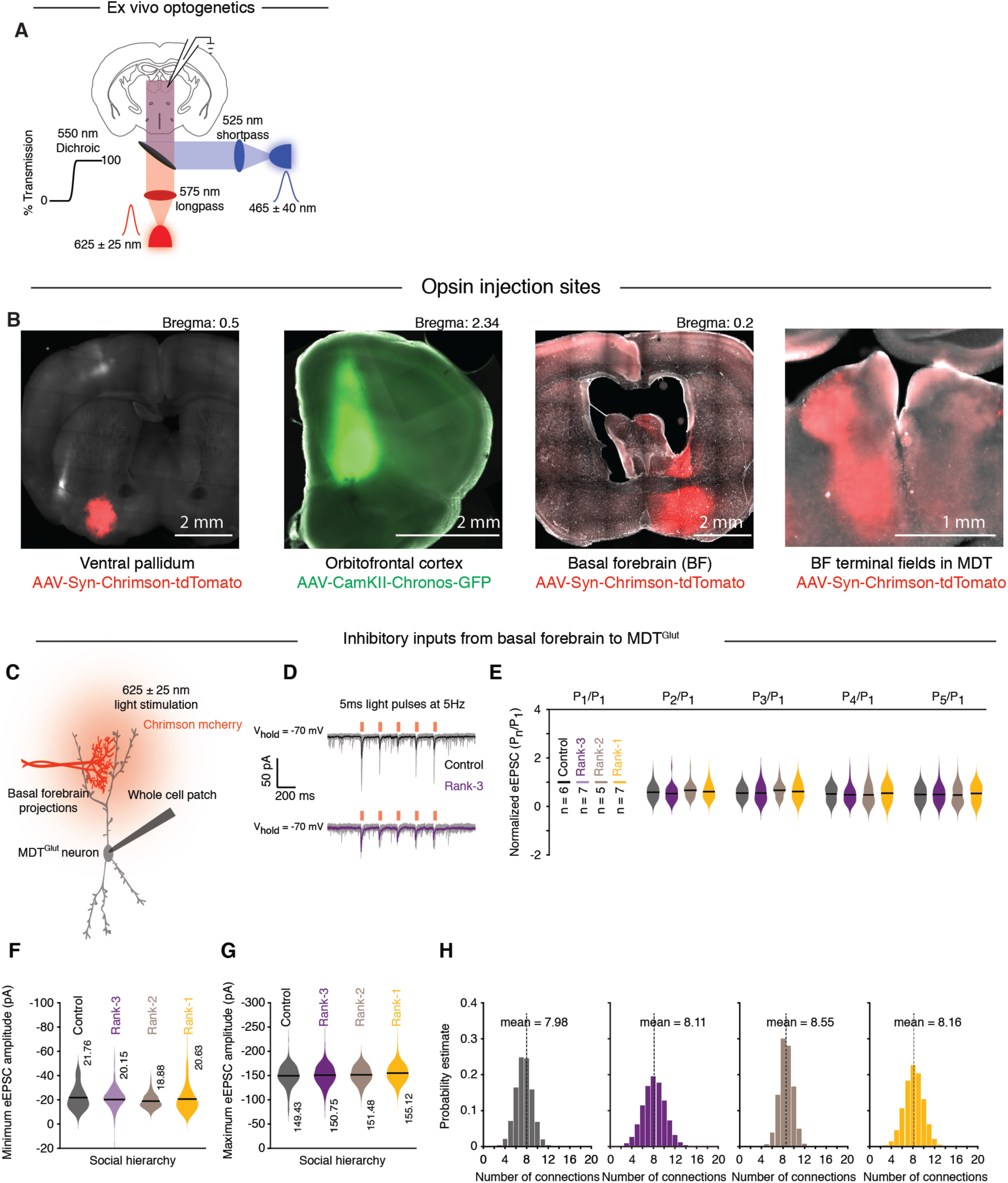
Inputs to medial dorsal thalamus. **(A**) Experimental set-up for activating optical actuators Chronos (with 465 nm light) and Chrimson (with 625 nm light). **(B)** Example of opsin injection sites in different brain areas ventral pallidum (AAV Chrimson, red, left), orbitofrontal cortex (AAV Chronos, green), basal forebrain (Chrimson, red) and projections of basal forebrain in the MDT (right). **(C)** Schematic depicting electrophysiological recordings of neurons in medial dorsal thalamus (MDT) slices in response to optogenetic activation of basal forebrain projections (BF, labelled with AAV Chrimson). **(D)** Top: electrophysiological traces (grey: 5 trials, black: mean response of 10 consecutive trials) for paired pulse depression for excitatory post synaptic activity (EPSCs) in response to activation of basal forebrain projections (light pulses: 5 ms duration, 625 nm at 5 Hz). Bottom: same paradigm results in similar paired pulse depression in rank-3 (purple) animals. **(E)** Summary plots depicting that the paired pulse paradigm results in depression in all ranks and control animals. **(F)** Minimum and **(G)** maximum evoked excitatory responses in MDT neurons in response to optogenetic activation of basal forebrain projections. **(H)** Histograms showing the distribution of number of excitatory synaptic connections between basal forebrain projections and a single MDT neuron for control (black), rank-1 (yellow), rank-2 (brown) and rank-3 (purple).

## SOM EXPERIMENTAL PROCEDURES AND METHODS

### Animals

Four mouse lines were used. C57BL/6J were shipped as littermates from Jackson laboratory. Vglut2-ires-Cre (Vong et al., 2011) (B. Lowell, Harvard Medical School) and PV-IRES-Cre (Hippenmeyer et al., 2005) (S. Arber, Friedrich Miescher Institute; gift from N. Uchida, Harvard University) mice were imported and bred in a barrier facility at Harvard University. PV-reporter mice (PV-IRES-Cre; Ai14-lsl-tdTomato) were a gift from T. Hensch (Harvard University). Animals were weaned at three weeks of age and maintained on a 12:12 hr light:dark cycle (lighted hours 02:00 to 14:00) with food and water ad libitum. Animal care and experiments were carried out in accordance with the NIH guidelines and approved by the Harvard University Institutional Animal Care and Use Committee (IACUC).

### Behavioral assays

All behavioral assays were performed on adult male mice (age > eight weeks) unless otherwise noted.

#### Social hierarchy paradigm and Elo ratings

Hierarchies were established in groups of three or four mice. Groups of four mice were used in assays testing hierarchy dynamics and individual performance. Groups of thee mice were used to assess the activity of MDT-ACC projection neurons and for electrophysiological recordings of the MDT. Groups were established by randomly picking one mouse from separate litters and matched for age (within two weeks) and weight (on average within 2 grams) and individually housed for on average five days before experiments began. Single housing promotes territorial behavior and allowed us to control for litter effects (e.g., maternal effects) and prior experiences among group members. Mice competed in tournaments of resident-intruder interactions and tube tests (Takahashi and Miczek, 2014). Each tournament was comprised of two round robins: the first consisted of each pairwise combination of animals, and the second consisted of the same pairwise combinations but with reversed ordering of the individuals (Fig. 1A). The group was first subjected to a tournament of resident-intruder interactions under dim red lighting. In groups of four, each tournament consisted of 12 interactions, where each mouse served as the resident and intruder in every unique pairing within a group of four mice. Each interaction lasted 30 minutes to 1 hour. The full tournament took on average six days. Next, the group was subjected to tournaments of the tube-test (Lindzey et al., 1961; Wang et al., 2011) under dim white light from floor lamps. Tournaments were repeated for at least seven days. Wins, losses, and time to decision were recorded.

Elo rating continuously updates player rating based on whether performance is better or worse than expected from previous contests; expected outcomes lead to smaller point changes than unexpected outcomes (Methods). Probabilistic models of dominance like Elo ratings provide an advantage over strict ordinal ranks because they can accommodate players tied for number of wins, quantify the magnitude of dominance between ranks, and are amenable to parametric statistics controlling for multiple response variables (Boyd and Silk, 1983). Elo ratings were determined with the EloRating R package (Neumann et al., 2011). Hierarchy stability was determined by applying the EloRating stability function as a moving average across each round robin. A stability of one indicates no rank changes—whereas a stability of zero indicates rank changes for all four animals—from one round robin to the next. Hierarchy consistency was determined for each unique pair of animals in a group and is a measurement of whether the outcome in one interaction is consistent with the next interaction. A sliding window of four interactions (in groups of four) or three interactions (in groups of three) determined consistency, where the count was reset to zero each time the outcome between a pair changed and reached a maximum of four for each window. Individual performance was determined with “delta Elo,” or the change in Elo from one round robin to the next.

For hierarchies receiving experimental manipulations (odor swab, MDT lesion, or CNO injection), manipulations were delivered only after the hierarchy reached stability criteria. Specifically, the hierarchy had to have a stable hierarchy of 1 for at least four round robins before the manipulation was made. For CNO injections, animals were habituated to mock i.p. injections prior to tests.

#### Defensive and aggressive behavior

Resident intruder interactions were video recorded in parallel using a multi-camera surveillance system (Geovision GV-1480 capture card) and CCD cameras. We defined defensive behavior as fleeing or avoiding the other mouse, including curling up in a C shape when being pursued or standing up on two feet and pushing away a pursuer mouse. Defensive frequency, duration, rate, and index (defined as fraction of temporal and count measurements of a given mouse during a pairwise interaction) were recorded using Observer XT 8 software (Noldus Information Technology).

#### Odor transfer experiments

Urine was collected by holding the mouse over a clean plexiglass sheet and urine was pipetted into a microfuge tube. Saliva was collected by pilocarpin injections under general anesthesia. Mice were anesthetized with 100 mg/kg ketamine (KetaVed, Vedco) and 10 mg/kg xylazine (AnaSed) via intra-peritoneal (i.p.) injection, and then i.p. injected with 0.5 mg/kg pilocarpine. Saliva was collected by pipetting directly from the mouth into a microfuge tube. In transfer experiments urine and saliva were either used fresh or frozen at −80**°**C and thawed immediately prior to the experiment. During odor transfer experiments, all mice in a social group were lightly anesthetized using isoflurane (VetOne) and washed with soap and water and padded down with 70% ethanol. Urine, saliva, or both was painted on to the face and L per substance). The painted animal was allowed to habituate to μ the swabbed odors for at least 10 minutes before beginning round robin tube test tournaments.

#### Sociability and declarative social memory assay

The nonconditioned social discrimination procedure was performed under dim red light essentially as described (Engelmann et al., 2011). Mice were habituated to the three-chamber apparatus (25.5 x 36 cm) consisting of two side chambers and a connecting corridor on multiple days before the test date. Sociability was measured during the learning phase of the experiment, where a juvenile (aged 16-35 days) was positioned under an inverted wire pencil cup in one side chamber and an empty inverted wire pencil cup was placed in the other chamber; the adult test animal was allowed to explore the apparatus for seven minutes. After seven minutes, test mice were given a 90-minute inter-trial intermission period in their home cage. Next, declarative memory was tested for 10 minutes during the recall phase, where the familiar juvenile from the learning phase was positioned under the same inverted pencil cup, and a novel juvenile was positioned on the opposite side under the pencil cup. The time spent in each of the three chambers was then measured and analyzed using Ethovision XT 8 software (Noldus Information Technology). In DREADD experiments, CNO or vehicle was delivered 30 minutes prior to the onset of the sociability assay.

#### Open field assay

Open field assay was used to measure locomotion and time spent in an exposed area of a cage. Mice were placed into a large cage (25.5 x 36 cm) with a clear acrylic lid for 15 minutes. Distance travelled, velocity, and time spent in the center of the cage (12 x 18 cm) was measured with Ethovision XT 8 software (Noldus Information Technology).

### Physiological measurements

Sperm and salivary glands were recovered immediately following sacrifice in mice that had established hierarchy.

#### Sperm count

*Cauda epydidymus* and *vas deferens* were bilaterally dissected and placed into 150 μL of PBS in a petri dish. For 12 minutes sperm were allowed to swim out, after which both organs were gently squeezed with forceps to press the remaining sperm out. All floating sperm were then collected by pipette and transferred into a microcentrifuge tube. To count sperm, 10 μL of the 150 μL suspension was mixed with 10 μL of Trypan Blue and cell numbers determined with Countess Automated Cell Counter (Thermo Fisher Scientific, C10227).

#### Salivary gland mass

Salivary glands, including both submaxillary and sublingual glands but not preputial gland, were bilaterally dissected as described (Jonjic, 2001) and individually weighed.

### Histology

All histological sections were mounted onto Superfrost slides (VWR, 48311-703) with DAPI-Vectashield (Vector Labs, H-1200). Sections were imaged at 10x resolution with an AxioScan Z1 slide scanner (Zeiss) or at 20x resolution in confocal stacks with an LSM 880 with Airyscan (Zeiss).

#### RNA in situ hybridization

In situ hybridization of *Cfos*, *Arc*, *Slc17a6* was performed as described (Wu et al., 2014). Fresh brain tissues were collected from animals in their home cage or 35 minutes after the mid-point of a tube test tournament, embedded in OCT (Tissue-Tek, 4583), and slowly frozen with dry ice. Whole brain, 20 μM coronal sections were kept at −80**°**C. Complementary DNA (cDNA) was cloned into approximately 800-base-pair segments into pCRII-TOPO vector (Thermo Fisher, K465040). Antisense complementary RNA probes were synthesized with T7 (Promega, P2075) or Sp6 polymerases (Promega, P1085) and labeled with Digoxigenin (DIG, Roche) or Fluorescein (FITC, Roche). Probes were hybridized at 68**°**C at a concentration of 0.5 to 1.0 ng/mL. Horseradish peroxidase conjugated antibodies against DIG (anti-DIG-POD, Roche, 1:500) and FITC (anti-FITC-POD, Roche, 1:250) incubated overnight and developed with the TSA-plus Cy3 system (Perkin Elmer). Signals were amplified with biotin-conjugated tyramide (Perkin Elmer) and developed with Alexa Fluor 488-conjugated streptavidin (Thermo Fisher, S11223). Washes were done with 0.5% PBST (0.5% Triton X-100 in PBS).

RNAscope in situ hybridization was performed by collecting frozen coronal sections (16 μM) as described above. Hybridizations were preformed using the RNAscope Multiplex Fluorescent Assay (ACDBio) according to the manufacturer’s instructions. ACDBio probes used were *Necab1* (#428541-C2), *Scn3a* (#502641), *Trpm3* (#459911) and *Cfos* (506921-C2, 506931-C2). Probes were visualized with ACDBio AMP 4 Alt B fluorescent dyes (C1 = Atto 550; C2 = Atto 550, C3 = Atto 647). Images were acquired on a Zeiss Axioscan microscope.

#### Fluorescence immunohistochemistry

Staining with CFOS, NeuN, GFAP and PV antibodies was performed according to standard protocols. Animals were perfused with phosphate buffered saline (PBS) followed by 4% paraformaldehyde (PFA). Dissected brain tissue was post-fixed in 4% PFA overnight and washed and stored in PBS. Whole brain, 60 µm sections were collected by embedding the brain in 4% low melting point agarose (Thermo Fisher Scientific, 16520-050) and sliced on a Vibratome (Leica). Sections were made permeable and blocked overnight against nonspecific fluorescence with blocking buffer (0.5% Triton X-100, 1% BSA, 2% normal donkey serum in PBS). Primary and secondary antibody incubations were in blocking buffer on a rotator at 6 hours room temperature or overnight at 4°C. Primary and secondary antibodies were washed with 0.5% PBST (5 x 60 min). Primary antibodies: rabbit anti-CFOS 1:2500 (Synaptic Systems 226003); mouse anti-NeuN 1:500 (Chemicon A60); rabbit anti-GFAP 1:000 (Abcam, Ab7260); rabbit anti-parvalbumin 1:1000 (Swant, PV27). Secondary antibodies (all from Thermo Fisher): Alexa-568 anti-goat (A-11057) 1:1,500, Alexa-555 anti-goat (A-21432) 1:1,500, and Alexa-647 anti-goat (A-21447) 1:1,500. CFOS antibody signal was amplified by incubation with Biotinylated horse anti-rabbit (Vector Laboratories, BA-1100) at 6 hours room temperature or overnight at 4°C and developed with Streptavidin, Alexa Fluor 555 Conjugate (Life Technologies, S32355); both steps were washed in PBST (5 x 60 min).

### Surgery

All surgeries were performed under aseptic conditions. Animals were anesthetized with 100 mg/kg ketamine (KetaVed, Vedco) and 10 mg/kg xylazine (AnaSed) via intra-peritoneal (i.p.) injection. Viruses and NMDA were stereotactically delivered with Nanoject II injector (Drummond Scientific) at a final volume of 135 nL per injection site. Analgesia (buprenorphine, 0.1 mg/kg, i.p.) was administered for 2 days following each surgery.

### Viruses

#### In vivo chemogenetic inhibition and excitation of MDT and cACC

Excitatory (pAAV-hSyn-DIO-hM3D(Gq)-mCherry) and inhibitory (pAAV-hSyn-DIO-hM4D(Gi)-mCherry) Serotype 8 DREADD-expressing viruses (Krashes et al., 2011) were produced from UNC Vector Core (discontinued) and Addgene (44361 and 44362).

#### Anterograde tracing

Anterograde tracing was performed in adult naïve Vglut2-ires-Cre male mice. MDT coordinates: AP: −1.35, ML: 0.43, DV: −3.2. A 1:1 mixture of AAV1/CAG-FLEx-tdTomato and AAV1/CAG-FLEx-Syn-GFP (Esposito et al., 2014) was injected to visualize presynaptic terminals of MPOA^Gal^ neurons.

#### Monosynaptic retrograde tracing

Input tracing experiments were performed in adult naïve Vglut2-ires-Cre and PV-IRES-Cre mice (cACC coordinates: AP: +1.18, ML: 0.17, DV: −1.3 to - 0.95) mice. Cre-dependent helper AAVs (Miyamichi et al., 2013) containing the TVA receptor for avian virus envelope protein (EnvA) and rabies glycoprotein (AAV-DIO-TVA-mCherry, AAV-DIO-RG) were injected in a 1:1 mixture. After 10 days of expression, EnvA-pseudotyped, G-deleted rabies virus (SBPN-RbV-EnvA-GFP, Salk Vector Core) (Wickersham et al. 2007) was injected into the same coordinates. After 4 days of expression, brains were prepared for histology.

#### *In vitro* optogenetic stimulation

Two channel-rhodopsin (ChR) variants under the human synapsin promoter (Syn) were used: ChrimsonR (AAV5-Syn-ChrimsonR-tdTomato-WPRE-bGH; V5453) and Chronos (AAV5-Syn-Chronos-GFP), both produced from the UNC Vector Core.

One variant with the CamKII promoter was used (AAV-CaMKII-Chronos-GFP, serotype AAV5), produced from the Duke Vector Core.

#### CTB tracing

Retrograde tracing experiments were performed in groups of C57BL/6J mice with established tube test hierarchies (among groups of three). Cholera toxin B (CTB) conjugated to Alexa Fluor-488, −555, or −647 (Thermo Fisher: CTB-488 C22841; CTB-555 C34776; CTB-647 C34778) was injected into the cACC in each mouse, as well as other areas known to receive projections from the MDT (dorsal-medial striatum, orbito-frontal cortex, agranular insular cortex, retrosplenial cortex). Five days after surgery, the mice were returned to daily tube test tournaments. Seven days after surgery, the mice were sacrificed 90 minutes after the mid-point of a tube test tournament and perfused with 4% PFA. Brains were post-fixed in 4% PFA, sectioned at 60 μm, and co-stained for CFOS. The number of CTB-CFOS co-labeled cells was quantified in the MDT.

#### MDT Lesions

N-methyl-D-aspartate (NMDA) in PBS (0.25 M) was bilaterally injected into MDT at a final volume 135 nL over a 20 min period. At the end of the injection, the pipet remained at the site for 5 min to allow for diffusion of the solution. Vehicle (PBS) was injected according to the same conditions. Lesions were confirmed with NeuN immunohistochemistry.

### Imaging and image analysis

Samples were imaged using an Axio Scan.Z1 slide scanner (Zeiss), and confocal stacks were acquired on an LSM 880 confocal microscope (Zeiss). Image processing was performed using custom routines for the Fiji distribution of ImageJ. Cell counts were performed by bilaterally cropping out anatomical regions of interest (ROI) followed by cellular segmentation with parameters customized to color channel and anatomical ROI.

For RNAscope puncta quantification, a custom FIJI script was used to manually crop left and right *Necab1*+ MDT brain regions; crops were separated into three separate images corresponding to DAPI (blue), *Necab1* (green) and the candidate gene (*Trpm3* or *Scn3a*) or *Cfos* in red. A CellProfiler pipeline (Kamentsky et al., 2011) was created to measure RNA signal of candidate genes within *Necab1*^+^ and *Necab1*^—^ cells. Nuclear boundaries (DAPI) were assigned by a global, minimum cross entropy algorithm. *Necab1* puncta were segmented using the adaptive Otsu thresholding method. Nuclear boundaries containing *Necab1* puncta were expanded by 4 microns to define the *Necab1*+ cells, whereas nuclear boundaries lacking *Necab1* puncta were defined as *Necab1*^—^ cells. To quantify puncta of candidate genes (i.e., *Trpm3* and *Scn3a*) within these two cell types, raw images were enhanced for speckle detection, segmented using the adaptive Otsu thresholding method, and puncta labeled with the RelateObjects function.

### Electrophysiology

#### Solutions

Modified ACSF (artificial cerebro-spinal fluid) contained (in mM): 105 choline chloride, 20 glucose, 24 NaHCO3, 2.5 KCl, 0.5 CaCl2, 8 MgSO4, 5 sodium ascorbate, 3 sodium pyruvate, 1.25 NaH2PO4 (osmolarity 290, pH 7.35) was used as cutting solution. All recordings were made in an oxygenated ACSF with composition (in mM): 115 NaCl, 2.5 KCl, 25 NaHCO3, 1.25 NaH2PO4, 1 MgSO4, 20 glucose, 2.0 CaCl2 (osmolarity 290, pH 7.35). The modified ACSF was used for isolating barium currents(in mM): 95 NaCl, 10 HEPES, 2.5 KCl, 1 MgCl2, 0.5 BaCl2, 20 glucose, 20 TEA, and 1.25 NaH2PO4, and 4 4-AP and 0.001 TTX.

Current-clamp internal solution contained (in mM): 120 potassium gluconate, 2.0 sodium gluconate, 10 HEPES, 4.0 Mg-ATP, 2.0 Na2-ATP, 0.3 Na3-GTP, and 4.0 NaCl (osmolarity 292, pH 7.34). Voltage clamp internal solution contained (in mM): 130 d-gluconic acid, 130 cesium hydroxide, 5.0 NaCl, 10 HEPES, 12 di-tris-phosphocreatine, 1 EGTA, 3.0 Mg-ATP, and 0.2 Na3-GTP (osmolarity 291, pH 7.35). To view cells after recording, internal solutions contained 1% biocytin or Alexa 488 or 594 as indicated. All chemicals were purchased from Sigma-Aldrich.

#### Acute brain slices

In vitro slice physiology was performed in adult (approximately two months) C57BL/6J mice with established tube test hierarchies (in groups of three) and in age-matched, individually housed controls. Mice were sacrificed during the inactive cycle at least two days after the last tube test tournament. Slices were prepared using methods described previously (Kapoor and Urban, 2006). Mice were lightly anesthetized with isoflurane exposure using a vaporizer (Datex-Ohmeda) connected to a clear acrylic chamber for two min, and then deeply anesthetized with an i.p. injection of a mixture of ketamine (100 mg/kg) and xylazine (10 mg/kg). Mice were transcardially perfused with ice-cold modified ACSF cutting solution, and brains were dissected into the same solution. Sagittal slices (300 µm thick) were obtained using a vibratome (VT1000S; Leica, Germany) and collected in ice-cold cutting solution. MDT was identified based on its proximity to anatomical landmarks including the fimbria, third ventricle, and mammillothlamic tract. cACC^PV^ neurons proximal to the genu of corpus callosum were identified by the tdTomato reporter. After cutting, slices were incubated in oxygenated ACSF solution at 35 °C for 45 min and then at room temperature for the duration of the experiment.

#### *In vitro* recordings

Whole-cell current-clamp and voltage-clamp recordings were made using borosilicate glass patch pipettes (6-10 Mohm) filled with current-clamp internal and voltage-clamp internal solutions, respectively, and slices maintained at 35 °C in oxygenated ACSF. Slices were visualized under custom-built infrared optics on a BX51WI microscope (Olympus Optical, Tokyo, Japan). Recordings were obtained with a Multiclamp 700B amplifier (Molecular Devices, Palo Alto, CA), and physiological data were collected via software written in LabView (National Instruments) and pClamp 10.3 (Molecular Devices). To visualize recorded cells, slices were fixed in 4% PFA overnight, washed with PBS, incubated in blocking buffer (0.3% Triton X-100, 2% normal goat serum in PBS) for four hours, stained overnight in blocking buffer with Streptavidin Alexa Fluor 594 or 647 conjugate (Life Technologies S11227) at 4 °C, and washed in 0.5% PBST. All neurons were multipolar as previously described in rodent MDT (Kuramoto et al., 2017).

For optogenetic axon terminal photostimulation experiments, Chronos or ChimsonR were activated using custom-built 465 nm (CBT-90-B-L11, Luminus) or 625 nm LEDs (CBT-90-RX-L15, Luminus). To record synaptic input to MDT cells, ChRs were transduced into cells of the following areas: orbitofrontal cortex (ORB), piriform cortex, lateral preoptic area, ventral pallidum, olfactory tubercle, and the diagonal band nucleus (HDB). Light pulses of 5ms, delivered at 1-10 Hz, were used to evoke synaptic transmission.

#### Short term plasticity measurements and simulation of number of input connections

Short term plasticity of inputs to MDT neurons was measured by measuring eEPSC amplitude of five successive evoked responses (10 ms light stimulus) separated by a 100 ms interval. Effects of social rank on the probability of release (e.g., facilitation or depression) were determined by dividing the current amplitude of each evoked response (i.e. first through fifth) by the amplitude of the first response.

To determine the minimal input of ChR expressing afferents (a proxy for a single synapse), we incremented light intensity starting at 0.5 mW/mm^2^ while holding MDT neurons at −70 mV, until 50% of light stimuli resulted in synaptic response. To determine the maximal input of ChR expressing afferents (a proxy for the total synaptic input), we incremented light intensity, until the synaptic response saturated. The relative number of inputs was then estimated using a simulation (Litvina and Chen, 2017), where a maximal response value was randomly chosen, and then individual minimal response values were randomly selected and summed until reaching the value of the maximum. This procedure was then repeated 100,000 times, and the distribution of the number of summed minimal responses was calculated. The mean number of minimal responses was used as the estimate for the relative number of connections.

#### TRPM3 pharmacology and barium current recordings

For pharmacological manipulation of TRPM3 channels, the agonist CIM 0216 (Tocris, 5521) and antagonist mefenamic acid (Sigma, 92574) were added separately to recording solution. MDT membrane voltage responses to the drugs were recorded in current clamp mode. Two experiments were done to address the role of axonal TRPM3 on synaptic input to PV neurons of the cACC. (1) mEPSCs in cACC^PV^ cells in response to mefenamic acid were recorded in voltage clamp mode in the presence of TTX. (2) Chronos was injected into the MDT and, at least three weeks later, light-evoked currents in cACC^PV^ cells were recorded in the presence of mefenamic acid.

Barium currents were recorded as a proxy for calcium currents to get better signal-to-noise ratio and to minimize inactivation (Dougalis et al., 2017; Hille, 2001), and were recorded in the presence of compounds to block sodium and potassium currents in modified oxygenated ACSF with composition (in mM): 95 NaCl, 2.5 KCl, 10 HEPES, 1.25 NaH2PO4, 1 MgCl2, 20 glucose, 0.5 BaCl2, 4 4-AP, 20 TEA and 1μm TTX (osmolarity 290, pH 7.35). Neurons were held −70 mV, hyperpolarized to −110 mV, and then depolarized in 5 mV increments to 30 mV. Current subtraction was used to isolate the TRPM3 component of the barium current.

#### Drugs

4-AP (Tocris, 0940); TTX (Hellobio, HB1034); CIM 0216 (selective TRPM3 channel agonist; Tocris, 5521). Mefenamic acid (blocker of Trpm3 channels; Sigma, 92574).

### Single nucleus RNA-Seq

#### Nuclear isolation, purification, and barcoding

Nuclei were extracted following a custom protocol designed to minimize the amount of time from euthanasia to nuclear barcoding (approximately four hours). Age-matched dominant, subordinate, and control mice from the same cohort were processed in parallel. Mouse brains were quickly dissected on ice and 2mm thick coronal sections were cut on a brain matrix (Zivic Instruments, #5325). Sections were placed on a chilled surface under a dissecting microscope and tissue punches from the thalamus were collected using a 2mm diameter cylindrical corer (Fine Science Tools, #18035-02). Tissue punches were pooled in chilled Hibernate-A Medium (ThermoFisher, #A1247501), minced into approximately 0.5^2^ mm pieces, and transferred to 3 mL lysis buffer (10 mM Tris-HCl, 10 mM NaCl, 3 mM MgCl2, 0.1% Nonidet P-40). Tissue was dounce homogenized in a glass tube 6-10 times with a grinder pestle (Potter-Elvehjem, #886000-0020) rotating at 300 rpm until opaque, transferred to a 15 mL tube, and pelleted in a swinging bucket centrifuge (Beckman Coulter GS-6R) at 50 RCF for 5 min at 4°C. Supernatant was removed and tissue carefully resuspended in 1.5 mL PBS with BSA (1%) and RNase inhibitor (0.2 U/μL, Promega, #N2615). To remove debris, suspension was passed over a 70 μM filter (ThermoFisher, #NC0120755) and then again over a 20 μM filter (ThermoFisher, #130-101-812). With the suspension on ice, nuclei were labeled by adding DAPI (1.5 μL, ThermoFisher, #D1306). DAPI-positive nuclei were sorted at 4°C on a FACSAria flow sorter (BD Immunocytometry Systems) with a 70 mm nozzle on the four-way high speed purity setting until approximately 100,000 nuclei were collected.

Finally, to estimate nuclear concentration, 10 uL of suspension was transferred to an Eppendorf tube and mixed with 1 μL propidium iodide (ThermoFisher, #P21493), counted on a Luna automated cell counter slide (Logos Biosystems), and adjusted to a final concentration of approximately 500 nuclei per microliter for 10X GEM formation. In total, 10 pools of nuclei were sequenced, where each pool consisted of thalamic microdissections from two mice (N = 20 mice). Nuclear pools were made from dominant (rank 1; 3 pools), subordinate (rank 3; 3 pools), and individually housed controls (4 pools).

#### Single nucleus library preparation and RNA-Seq

Using the 10X Genomics droplet microfluidics Gemcode platform (Zheng et al., 2017), nuclei from the suspension were loaded onto Chromium instrument. Libraries were generated using Chromium v2 reagents, and cDNA amplified according to the manufacturer’s instructions. Samples were sequenced on an Illumina Nextseq at the Harvard University Bauer Core Facility. We recovered an average of 4,699 transcripts and 2,169 unique genes per nucleus (SOM Fig. 6B).

#### Sequence alignment and identification of variable genes

Transcript sequences were demultiplexed, aligned to the mm10 mouse genome, counted, and assembled into gene-cell matrices using the 10x Genomics Cell Ranger pipeline with default parameters. The matrices were then analyzed in Seurat version 2.3.4 (Butler et al., 2018). Gene expression for each nucleus was log-normalized and scaled (i.e., log transformed, normalized by total transcript count, and multiplied by a factor of 10,000). To eliminate aberrant droplets, we filtered out nuclei based on mitochondrial content (> 10%), nuclear doublets (> 20,000 UMIs), red blood cells (>1% hemoglobin genes), and immediate early gene (IEG) responses (e.g., responses to the dissection/dissociation process; > 1% IEG content). Glial cells were filtered out based on the expression of established glial-specific genes. To identify variable genes that characterize cellular heterogeneity based on dispersion (variance/mean) in expression level, we used the FindVariableGenes function with the following parameters (x.low.cutoff = .1, y.cutoff = .8). To control for unwanted variation in nuclear expression profiles of the variable genes, we used regression to control for the effects of nuclear UMI count, percent mitochondria, percent IEG, and batch identity using the ScaleData function using ‘negbinom’ for the model parameter; the residual matrix was then scaled and centered and used for downstream analysis.

#### Dimensionality reduction, clustering, and marker gene identification

Dimensionality reduction was performed with principal components analysis (PCA) of the variable gene residual matrices as implemented in Seurat (Satija et al., 2015). To determine the number of principal components (PCs) to use in downstream analyses, we used the JackStraw function to compare PCA scores of random subsets of genes with the observed data, allowing us to find significant PCs with a strong enrichment of genes with low P-values. We also used the PCElbowPlot function to visualize the cumulative standard deviations of each PC and the point at which successive PCs explained a diminishing degree of variance.

We defined cell clusters by passing the first 45 PCs to a shared nearest neighbor (SNN) clustering algorithm as implemented in the FindClusters function; for this analysis we used the smart local moving (SLM) community detection algorithm and a K-nearest neighbor value of 19. To visualize nuclei according to their PC scores, we used the UMAP algorithm to place each nucleus on a two-dimensional plot; subsequently, each nucleus was colored based on cluster identity.

To nominate cluster-specific marker genes based on differential expression, we used the FindAllMarker function with the MAST algorithm (Finak et al., 2015); differentially expressed genes that were expressed in at least 15% of cells within the cluster and with a fold change of more than 0.5 (log scale) were selected as markers. To determine whether clusters corresponded to novel or well-established cell types (e.g. inhibitory, excitatory, oligodendrocytes), we examined the expression patterns in the mouse brain using the Allen Brain Atlas (Lein et al., 2006).

#### Thalamic sNuc-Seq and validation

We identified a distinct cluster corresponding to the reticular thalamus, a known inhibitory (GABAergic) population (Krol et al., 2018) that robustly expressed *Gad1*, *Gad2,* and the new marker *Isl1* (SOM Fig. 6C, E). The majority of sequenced nuclei (20,533, 61%) belonged to one large metacluster comprised of 13 clusters, each identified to be excitatory thalamic areas based on expression of *Vglut1* and *Vglut2* (SOM Fig. 6E) (Lein et al., 2006). Nine of these 13 clusters were assigned to paraventricular, dorsal, and ventral thalamic regions, and the remaining five assigned to the MDT.

### Statistical Analyses

We used linear models (lm), linear mixed-effect regression (lmer) models, generalized linear models (glm), and Wilcoxon-rank sum tests (Bates et al., 2015). Where appropriate, data were transformed to achieve more normal distributions. *Post hoc* analyses were done using ANOVA, Tukey’s honest significant difference (HSD) test, Dunn’s multiple comparisons, or least-square means contrasts. A value of p ≤ 0.05 was considered significant. All analyses were done in R (3.1.3) and Matlab.

#### Behavioral assays

To determine differences in hierarchy stability and pairwise consistency between observed and random data (generated by selecting a random outcome for an equivalent number of trials per group for that of the observed data), we used lmer. Stability and consistency were the response variables and explanatory variables were condition (observed vs. random), tube rank, and social group as a random effect.

To investigate the strength of association between resident-intruder defensive behaviors (i.e., defensive duration or defensive index) and tube rank, we used Akaike information criterion (AIC) selection of three nested models, each with tube rank as the response variable. In the full model, explanatory variables were defensive duration, defensive index, and social group as a random effect; in the second model, defensive duration was excluded; in the third model, defensive index was excluded. To determine the probability of defensive behavior based on tube rank, we used a binomial logistic regression by modeling defensiveness (yes or no) as the response variable and explanatory variables included simplified tube rank (upper rank vs. lower rank) and interaction time.

To investigate the role of the MDT ablation in hierarchy dynamics, we first used a sliding window to measure hierarchy stability and pairwise consistency on a per-round-robin basis. We next used lmer by modeling stability or consistency as the response variable and explanatory variables were condition (lesion or sham) and social group as a random effect. Pairwise contrasts were determined by Dunn’s multiple comparison test. To determine the effect of MDT ablation on decision timing in the tube test, we used a lmer with decision time as the response variable, and explanatory variables were condition and tournament number.

To investigate the role of the MDT in pheromonal regulation of social hierarchy, we determined the performance of rank4 males when swabbed with rank1 urine in the lesion and sham groups. A lmer was used to model delta Elo as the response variable and explanatory variables included swab condition (untreated, swab, or washed as a swab control), tube rank, the interaction between swab condition and tube rank, and social group as a random effect. Post hoc comparisons were made using least-square means contrasts.

#### Physiological measurements

To investigate the association between tube rank and testosterone-dependent traits we used lmer by modeling sperm count or salivary gland weight as the response variable and explanatory variables included tube rank, body mass, and social group as a random effect.

#### Histology, tracing, and cellular image analysis

To investigate the association between brain-wide neural activity and tube rank, we used lmer by modeling the percentage of *Cfos*-positive cells as the response variable and explanatory variables included tube rank and social group as a random effect. We first used the two-sided t-statistic to investigate correlation matrices of *Cfos* expression between each brain region in rank1 and rank4 males. Next, do determine the effect of rank on brainwide *Cfos* correlations, we evaluated the difference between the rank1 and rank4 t-statistic matrices.

To investigate the effect of tube rank on the activity of MDT-cACC projection neurons, a lmer was used to model the number of CFOS-CTB co-labeled cells as the response variable and explanatory variables included tube rank and social group as a random effect.

To investigate the effect of tube rank on the number of puncta within genetically define cell types, a LMER was used to test the number of puncta per cell type as the response variable; explanatory variables included cell area and tube rank as fixed effects, and social group and histological slice ID were included as random effects.

#### In vivo chemogenetic inhibition and excitation of MDT and cACC

To investigate the effect of chemogenetic neuronal manipulation on individual performance we modeled change in Elo as the response variable and explanatory variables were CNO condition (untreated, saline, or CNO), tube rank, the interaction between CNO condition and tube rank, and social group as a random effect. To determine the significance of CNO treatment on individual performance, *post hoc* comparisons were made using least-square means contrasts.

#### Electrophysiology recordings

Whole cell patch clamp data were analyzed as previously described (Kapoor et al., 2016). To investigate the boundary between two clusters of cells based on membrane resistance we used K-means clustering. Wilcoxon rank-sum tests were used to compare the distributions of cluster-1 and cluster-2, and to test the effect of rank on the frequency and size of EPSCs and IPSCs. We used non-parametric two sample Kolmogorov-Smirnov test (KS test) to calculate the significance of mefenamic acid on firing rate of MDT neurons. For comparisons of pharmacological effects of mefenamic acid on the spontaneous activity of PV neurons, the KS test was used. Similarly, KS test was used to compare the amplitude of evoked excitatory events in cACC^PV^ neuron.

To compare the paired pulse responses across animals we used Wilcoxon ranksum test. We also used Wilcoxon ranksum test to compare the amplitudes of minimum and maximum evoked EPSCs and IPSCs. Similarly, ranksum test was used to compare the number of connections between OFC and MDT neurons, and basal forebrain and MDT neurons across different ranks.

#### Effects of social status on cell type specific differential gene expression and gene set enrichment

Nuclei were classified according to their condition (dominant, subordinate, or control). Differentially expressed genes between each pairwise comparison (rank 1 vs. control; rank 2 vs. control; rank 1 vs. rank3) on a per-cluster basis were identified with the Seurat function FindAllMarkers using the MAST algorithm. For each cluster, only genes that were expressed in at least 5% of nuclei were considered.

To determine the effect of social status on gene set enrichment, we used gene set enrichment analysis (GSEA), a technique designed to detect modest but coordinate changes in the expression of groups of functionally related genes that are defined *a priori* (Mootha et al., 2003, 2004). We generated a custom list of 145 gene sets pertaining to neural structure and function (Appendix 1). Gene sets were derived from four sources as described previously (Henry et al., 2015) with some modifications. (1) The IUPHAR Database (Pawson et al., 2014), a curated list of genes encoding biological targets of small molecules and licensed drugs. (2) The PANTHER Mus musculus database (Mi et al., 2010), a database of gene families and their functionally related subfamilies used to classify gene product function. (3) Child terms under the Gene Ontology (GO; version 2018) parent term “Synapse,” (GO:0045202). (4) Child terms under the parent term “Sequence-specific DNA binding transcription factor activity” (GO: 0003700). To be included in GSEA, each gene set had to have between 15 and 500 genes.

Enrichment score was calculated from the signal-to-noise ratio of each gene for each pairwise comparison of log-normalized gene counts, and this metric was weighted to ensure that low expression, low variance genes did not contribute to a positive enrichment score. We then performed a permutation test for false discovery rate (FDR) by permuting condition assignments 1,000 times. Gene sets with an FDR < 0.25 and an adjusted P-value < 0.01 were considered significant. Leading-edge genes were defined as those genes appearing in the ranked list before the point at which the running sum recaches its maximum deviation from zero. Finally, to examine the intersection of GSEA and DEG datasets, we used a union of GSEA leading edge genes with differentially expressed genes exhibiting a log-fold change of at least 0.1 on a per cluster, per pairwise comparison basis.

## Zotero References

Alexander, G.M., Rogan, S.C., Abbas, A.I., Armbruster, B.N., Pei, Y., Allen, J.A., Nonneman, R.J., Hartmann, J., Moy, S.S., Nicolelis, M.A., et al. (2009). Remote control of neuronal activity in transgenic mice expressing evolved G protein-coupled receptors. Neuron 63, 27–39.

Armbruster, B.N., Li, X., Pausch, M.H., Herlitze, S., and Roth, B.L. (2007). Evolving the lock to fit the key to create a family of G protein-coupled receptors potently activated by an inert ligand. Proc Natl Acad Sci U A 104, 5163–5168.

Bargmann, C.I. (2012). Beyond the connectome: How neuromodulators shape neural circuits. Bioessays 34, 458–465.

Bates, D., Mächler, M., Bolker, B., and Walker, S. (2015). Fitting Linear Mixed-Effects Models Using lme4. J. Stat. Softw. Vol 1 Issue 1 2015.

Baulieu, E.E. (1998). NEUROSTEROIDS: A NOVEL FUNCTION OF THE BRAIN. Psychoneuroendocrinology 23, 963–987.

Beiderbeck, D.I., Neumann, I.D., and Veenema, A.H. (2007). Differences in intermale aggression are accompanied by opposite vasopressin release patterns within the septum in rats bred for low and high anxiety. Eur. J. Neurosci. 26, 3597–3605.

Boyd, R., and Silk, J.B. (1983). A method for assigning cardinal dominance ranks. Anim. Behav. 31, 45–58.

Bronson, F.H. (1979). The Reproductive Ecology of the House Mouse. Q. Rev. Biol. 54, 265–299.

Butler, A., Hoffman, P., Smibert, P., Papalexi, E., and Satija, R. (2018). Integrating single-cell transcriptomic data across different conditions, technologies, and species. Nat. Biotechnol. 36, 411.

Chakraborty, S., Kolling, N., Walton, M.E., and Mitchell, A.S. (2016). Critical role for the mediodorsal thalamus in permitting rapid reward-guided updating in stochastic reward environments. ELife 5, e13588.

Chiao, J.Y. (2010). Neural basis of social status hierarchy across species. Curr Opin Neurobiol 20, 803–809.

Chou, M.-Y., Amo, R., Kinoshita, M., Cherng, B.-W., Shimazaki, H., Agetsuma, M., Shiraki, T., Aoki, T., Takahoko, M., Yamazaki, M., et al. (2016). Social conflict resolution regulated by two dorsal habenular subregions in zebrafish. Science 352, 87.

Courtin, J., Chaudun, F., Rozeske, R.R., Karalis, N., Gonzalez-Campo, C., Wurtz, H., Abdi, A., Baufreton, J., Bienvenu, T.C.M., and Herry, C. (2013). Prefrontal parvalbumin interneurons shape neuronal activity to drive fear expression. Nature 505, 92.

Courtiol, E., and Wilson, D. (2015). The olfactory thalamus: unanswered questions about the role of the mediodorsal thalamic nucleus in olfaction. Front. Neural Circuits 9, 49.

Creel, S. (2001). Social dominance and stress hormones. Trends Ecol. Evol. 16, 491–497.

Creel, S., Dantzer, B., Goymann, W., and Rubenstein, D.R. (2012). The ecology of stress: effects of the social environment. Funct. Ecol. 27, 66–80.

Davis, K.D., Taylor, K.S., Hutchison, W.D., Dostrovsky, J.O., McAndrews, M.P., Richter, E.O., and Lozano, A.M. (2005). Human Anterior Cingulate Cortex Neurons Encode Cognitive and Emotional Demands. J. Neurosci. 25, 8402.

Delevich, K., Tucciarone, J., Huang, Z.J., and Li, B. (2015). The Mediodorsal Thalamus Drives Feedforward Inhibition in the Anterior Cingulate Cortex via Parvalbumin Interneurons. J. Neurosci. 35, 5743–5753.

Dougalis, A.G., Matthews, G.A.C., Liss, B., and Ungless, M.A. (2017). Ionic currents influencing spontaneous firing and pacemaker frequency in dopamine neurons of the ventrolateral periaqueductal gray and dorsal raphe nucleus (vlPAG/DRN): A voltage-clamp and computational modelling study. J. Comput. Neurosci. 42, 275–305.

Elo, A.E. (1978). The Rating of Chess Players, Past and Present (New York: Arco).

Engelmann, M., Hadicke, J., and Noack, J. (2011). Testing declarative memory in laboratory rats and mice using the nonconditioned social discrimination procedure. Nat Protoc. 6, 1152–1162.

Esposito, M.S., Capelli, P., and Arber, S. (2014). Brainstem nucleus MdV mediates skilled forelimb motor tasks. Nature 508, 351–356.

Fan, Z., Zhu, H., Zhou, T., Wang, S., Wu, Y., and Hu, H. (2019). Using the tube test to measure social hierarchy in mice. Nat. Protoc. 14, 819–831.

Ferguson, B.R., and Gao, W.-J. (2018). Thalamic Control of Cognition and Social Behavior Via Regulation of Gamma-Aminobutyric Acidergic Signaling and Excitation/Inhibition Balance in the Medial Prefrontal Cortex. Biol Psychiatry 83, 657–669.

Fillinger, C., Yalcin, I., Barrot, M., and Veinante, P. (2017). Afferents to anterior cingulate areas 24a and 24b and midcingulate areas 24a and 24b in the mouse. Brain Struct. Funct. 222, 1509–1532.

Fillinger, C., Yalcin, I., Barrot, M., and Veinante, P. (2018). Efferents of anterior cingulate areas 24a and 24b and midcingulate areas 24a and 24b in the mouse. Brain Struct. Funct. 223, 1747–1778.

Finak, G., McDavid, A., Yajima, M., Deng, J., Gersuk, V., Shalek, A.K., Slichter, C.K., Miller, H.W., McElrath, M.J., Prlic, M., et al. (2015). MAST: a flexible statistical framework for assessing transcriptional changes and characterizing heterogeneity in single-cell RNA sequencing data. Genome Biol. 16, 278.

Frankland, P.W., Bontempi, B., Talton, L.E., Kaczmarek, L., and Silva, A.J. (2004). The Involvement of the Anterior Cingulate Cortex in Remote Contextual Fear Memory. Science 304, 881–883.

Franklin, T.B., Silva, B.A., Perova, Z., Marrone, L., Masferrer, M.E., Zhan, Y., Kaplan, A., Greetham, L., Verrechia, V., Halman, A., et al. (2017). Prefrontal cortical control of a brainstem social behavior circuit. Nat. Neurosci. 20, 260.

Frans B. M. de Waal (1986). The Integration of Dominance and Social Bonding in Primates. Q. Rev. Biol. 61, 459–479.

Ghazanfar, A.A., and Santos, L.R. (2004). Primate brains in the wild: the sensory bases for social interactions. Nat. Rev. Neurosci. 5, 603–616.

Golden, S.A., Heshmati, M., Flanigan, M., Christoffel, D.J., Guise, K., Pfau, M.L., Aleyasin, H., Menard, C., Zhang, H., Hodes, G.E., et al. (2016). Basal forebrain projections to the lateral habenula modulate aggression reward. Nature 534, 688.

Gröschl, M. (2009). The physiological role of hormones in saliva. BioEssays 31, 843–852.

Guhl, A.M. (1956). The social order of chickens. Sci. Am. 194, 42–46.

Halassa, M.M., and Kastner, S. (2017). Thalamic functions in distributed cognitive control. Nat. Neurosci. 20, 1669–1679.

Hawley, P.H. (1999). The Ontogenesis of Social Dominance: A Strategy-Based Evolutionary Perspective. Dev. Rev. 19, 97–132.

Held, K., Voets, T., and Vriens, J. (2015). TRPM3 in temperature sensing and beyond. Temp. Austin Tex 2, 201–213.

Held, K., Gruss, F., Aloi, V.D., Janssens, A., Ulens, C., Voets, T., and Vriens, J. (2018). Mutations in the voltage-sensing domain affect the alternative ion permeation pathway in the TRPM3 channel. J. Physiol. 596, 2413–2432.

Henry, F.E., Sugino, K., Tozer, A., Branco, T., and Sternson, S.M. (2015). Cell type-specific transcriptomics of hypothalamic energy-sensing neuron responses to weight-loss. Elife Camb. 4, e09800.

Hille, B. (2001). Ionic Channels of Excitable Membranes (Sinauer Associates, Inc.).

Hippenmeyer, S., Vrieseling, E., Sigrist, M., Portmann, T., Laengle, C., Ladle, D.R., and Arber, S. (2005). A Developmental Switch in the Response of DRG Neurons to ETS Transcription Factor Signaling. PLoS Biol 3, e159.

Hitti, F.L., and Siegelbaum, S.A. (2014). The hippocampal CA2 region is essential for social memory. Nature 508, 88–92.

Huchard, E., English, S., Bell, M.B.V., Thavarajah, N., and Clutton-Brock, T. (2016). Competitive growth in a cooperative mammal. Nature 533, 532.

Hurst, J.L., Beynon, R.J., Humphries, R., Malone, N., Nevison, C.M., Payne, C., Robertson, D.H., and Veggerby, C. (2001). Information in Scent Signals of Competitive Social Status: The Interface Between Behaviour and Chemistry. In Chemical Signals in Vertebrates, A. Marchlewska-Koj, J.J. Lepri, and D. Müller-Schwarze, eds. (Boston, MA: Springer), p.

Jackman, S.L., Turecek, J., Belinsky, J.E., and Regehr, W.G. (2016). The calcium sensor synaptotagmin 7 is required for synaptic facilitation. Nature 529, 88.

Jonjic, S. (2001). Surgical Removal of Mouse Salivary Glands. In Current Protocols in Immunology, (John Wiley & Sons, Inc.), p.

Kamentsky, L., Jones, T.R., Fraser, A., Bray, M.-A., Logan, D.J., Madden, K.L., Ljosa, V., Rueden, C., Eliceiri, K.W., and Carpenter, A.E. (2011). Improved structure, function and compatibility for CellProfiler: modular high-throughput image analysis software. Bioinformatics 27, 1179–1180.

Kapoor, V., and Urban, N.N. (2006). Glomerulus-Specific, Long-Latency Activity in the Olfactory Bulb Granule Cell Network. J. Neurosci. 26, 11709.

Kapoor, V., Provost, A.C., Agarwal, P., and Murthy, V.N. (2016). Activation of raphe nuclei triggers rapid and distinct effects on parallel olfactory bulb output channels. Nat. Neurosci. 19, 271–282.

Klose, C., Straub, I., Riehle, M., Ranta, F., Krautwurst, D., Ullrich, S., Meyerhof, W., and Harteneck, C. (2011). Fenamates as TRP channel blockers: mefenamic acid selectively blocks TRPM3. Br. J. Pharmacol. 162, 1757–1769.

Krashes, M.J., Koda, S., Ye, C., Rogan, S.C., Adams, A.C., Cusher, D.S., Maratos-Flier, E., Roth, B.L., and Lowell, B.B. (2011). Rapid, reversible activation of AgRP neurons drives feeding behavior in mice. J Clin Invest 121, 1424–1428.

Krettek, J.E., and Price, J.L. (1977). The cortical projections of the mediodorsal nucleus and adjacent thalamic nuclei in the rat. J. Comp. Neurol. 171, 157–191.

Krol, A., Wimmer, R.D., Halassa, M.M., and Feng, G. (2018). Thalamic Reticular Dysfunction as a Circuit Endophenotype in Neurodevelopmental Disorders. Neuron 98, 282–295.

Kuramoto, E., Pan, S., Furuta, T., Tanaka, Y.R., Iwai, H., Yamanaka, A., Ohno, S., Kaneko, T., Goto, T., and Hioki, H. (2017). Individual mediodorsal thalamic neurons project to multiple areas of the rat prefrontal cortex: A single neuron-tracing study using virus vectors. J. Comp. Neurol. 525, 166–185.

Kuroda, M., López-Mascaraque, L., and Price, J. (1992). Neuronal and synaptic composition of the mediodorsal thalamic nucleus in the rat: A light and electron microscopic golgi study. J. Comp. Neurol. 326, 61–81.

Kuroda, M., Yokofujita, J., and Murakami, K. (1998). An ultrastructural study of the neural circuit between the prefrontal cortex and the mediodorsal nucleus of the thalamus. Prog. Neurobiol. 54, 417–458.

Kvitsiani, D., Ranade, S., Hangya, B., Taniguchi, H., Huang, J.Z., and Kepecs, A. (2013). Distinct behavioural and network correlates of two interneuron types in prefrontal cortex. Nature 498, 363–366.

Lein, E.S., Hawrylycz, M.J., Ao, N., Ayres, M., Bensinger, A., Bernard, A., Boe, A.F., Boguski, M.S., Brockway, K.S., Byrnes, E.J., et al. (2006). Genome-wide atlas of gene expression in the adult mouse brain. Nature 445, 168.

Ligneul, R., Obeso, I., Ruff, C.C., and Dreher, J.-C. (2016). Dynamical Representation of Dominance Relationships in the Human Rostromedial Prefrontal Cortex. Curr. Biol. 26, 3107–3115.

Lin, D., Boyle, M.P., Dollar, P., Lee, H., Lein, E.S., Perona, P., and Anderson, D.J. (2011). Functional identification of an aggression locus in the mouse hypothalamus. Nature 470, 221.

Lindzey, G., Winston, H., and Manosevitz, M. (1961). Social Dominance in Inbred Mouse Strains. Nature 191, 474–476.

Litvina, E.Y., and Chen, C. (2017). Functional Convergence at the Retinogeniculate Synapse. Neuron 96, 330–338.e5.

Lu, J., Tucciarone, J., Padilla-Coreano, N., He, M., Gordon, J.A., and Huang, Z.J. (2017). Selective inhibitory control of pyramidal neuron ensembles and cortical subnetworks by chandelier cells. Nat. Neurosci. 20, 1377.

Macosko, E.Z., Basu, A., Satija, R., Nemesh, J., Shekhar, K., Goldman, M., Tirosh, I., Bialas, A.R., Kamitaki, N., Martersteck, E.M., et al. (2015). Highly Parallel Genome-wide Expression Profiling of Individual Cells Using Nanoliter Droplets. Cell 161, 1202–1214.

Marder, E., O’Leary, T., and Shruti, S. (2014). Neuromodulation of Circuits with Variable Parameters: Single Neurons and Small Circuits Reveal Principles of State-Dependent and Robust Neuromodulation. Annu. Rev. Neurosci. 37, 329–346.

Maren, S., Aharonov, G., and Fanselow, M.S. (1997). Neurotoxic lesions of the dorsal hippocampus and Pavlovian fear conditioning in rats. Behav. Brain Res. 88, 261–274.

McHenry, J.A., Otis, J.M., Rossi, M.A., Robinson, J.E., Kosyk, O., Miller, N.W., McElligott, Z.A., Budygin, E.A., Rubinow, D.R., and Stuber, G.D. (2017). Hormonal gain control of a medial preoptic area social reward circuit. Nat. Neurosci. 20, 449.

Mi, H., Dong, Q., Muruganujan, A., Gaudet, P., Lewis, S., and Thomas, P.D. (2010). PANTHER version 7: improved phylogenetic trees, orthologs and collaboration with the Gene Ontology Consortium. Nucleic Acids Res. 38, D204–D210.

Miczek, K.A., Maxson, S.C., Fish, E.W., and Faccidomo, S. (2001). Aggressive behavioral phenotypes in mice. Behav Brain Res 125, 167–181.

Miller, O.H., Bruns, A., Ben Ammar, I., Mueggler, T., and Hall, B.J. (2017). Synaptic Regulation of a Thalamocortical Circuit Controls Depression-Related Behavior. Cell Rep. 20, 1867–1880.

Miyamichi, K., Shlomai-Fuchs, Y., Shu, M., Weissbourd, B.C., Luo, L., and Mizrahi, A. (2013). Dissecting Local Circuits: Parvalbumin Interneurons Underlie Broad Feedback Control of Olfactory Bulb Output. Neuron 80, 1232–1245.

Moffitt, J.R., Bambah-Mukku, D., Eichhorn, S.W., Vaughn, E., Shekhar, K., Perez, J.D., Rubinstein, N.D., Hao, J., Regev, A., Dulac, C., et al. (2018). Molecular, spatial, and functional single-cell profiling of the hypothalamic preoptic region. Science 362, eaau5324.

Mootha, V.K., Lindgren, C.M., Eriksson, K.-F., Subramanian, A., Sihag, S., Lehar, J., Puigserver, P., Carlsson, E., Ridderstråle, M., Laurila, E., et al. (2003). PGC-1α-responsive genes involved in oxidative phosphorylation are coordinately downregulated in human diabetes. Nat. Genet. 34, 267.

Mootha, V.K., Daly, M.J., Patterson, N., Hirschhorn, J.N., Groop, L.C., and Altshuler, D. (2004). Reply to “Statistical concerns about the GSEA procedure.” Nat. Genet. 36, 663.

Nelson, R.J., and Trainor, B.C. (2007). Neural mechanisms of aggression. Nat. Rev. Neurosci. 8, 536.

Nelson, A.C., Cunningham, C.B., Ruff, J.S., and Potts, W.K. (2015). Protein pheromone expression levels predict and respond to the formation of social dominance networks. J. Evol. Biol. 28, 1213–1224.

Neumann, C., Duboscq, J., Dubuc, C., Ginting, A., Irwan, A.M., Agil, M., Widdig, A., and Engelhardt, A. (2011). Assessing dominance hierarchies: validation and advantages of progressive evaluation with Elo-rating. Anim. Behav. 82, 911–921.

Noonan, M.P., Sallet, J., Mars, R.B., Neubert, F.X., O’Reilly, J.X., Andersson, J.L., Mitchell, A.S., Bell, A.H., Miller, K.L., and Rushworth, M.F.S. (2014). A Neural Circuit Covarying with Social Hierarchy in Macaques. PLoS Biol 12, e1001940.

Oberwinkler, J., Lis, A., Giehl, K.M., Flockerzi, V., and Philipp, S.E. (2005). Alternative Splicing Switches the Divalent Cation Selectivity of TRPM3 Channels. J. Biol. Chem. 280, 22540–22548.

Parnaudeau, S., Taylor, K., Bolkan, S.S., Ward, R.D., Balsam, P.D., and Kellendonk, C. (2015). Mediodorsal Thalamus Hypofunction Impairs Flexible Goal-Directed Behavior. Biol. Psychiatry 77, 445–453.

Pawson, A.J., Sharman, J.L., Benson, H.E., Faccenda, E., Alexander, S.P.H., Buneman, O.P., Davenport, A.P., McGrath, J.C., Peters, J.A., Southan, C., et al. (2014). The IUPHAR/BPS Guide to PHARMACOLOGY: an expert-driven knowledgebase of drug targets and their ligands. Nucleic Acids Res 42, D1098–D1106.

Phillips, J.W., Schulmann, A., Hara, E., Winnubst, J., Liu, C., Valakh, V., Wang, L., Shields, B.C., Korff, W., Chandrashekar, J., et al. (2019). A repeated molecular architecture across thalamic pathways. Nat. Neurosci.

Price, J.L., and Slotnick, B.M. (1983). Dual olfactory representation in the rat thalamus: An anatomical and electrophysiological study. J. Comp. Neurol. 215, 63–77.

Qu, C., Ligneul, R., Van der Henst, J.-B., and Dreher, J.-C. (2017). An Integrative Interdisciplinary Perspective on Social Dominance Hierarchies. Trends Cogn. Sci. 21, 893–908.

Regehr, W.G. (2012). Short-Term Presynaptic Plasticity. Cold Spring Harb. Perspect. Biol. 4.

Rosner, W., Auchus, R.J., Azziz, R., Sluss, P.M., and Raff, H. (2007). Utility, Limitations, and Pitfalls in Measuring Testosterone: An Endocrine Society Position Statement. J. Clin. Endocrinol. Metab. 92, 405–413.

Sapolsky, R.M. (2004). Social Status and Health in Humans and Other Animals. Annu. Rev. Anthropol. 33, 393–418.

Sapolsky, R.M. (2005). The influence of social hierarchy on primate health. Science 308, 648–652.

Satija, R., Farrell, J.A., Gennert, D., Schier, A.F., and Regev, A. (2015). Spatial reconstruction of single-cell gene expression data. Nat. Biotechnol. 33, 495.

Sawada, K., and Noumura, T. (1991). Effects of Castration and Sex Steroids on Sexually Dimorphic Development of the Mouse Submandibular Gland. Cells Tissues Organs 140, 97–103.

Schmitt, L.I., Wimmer, R.D., Nakajima, M., Happ, M., Mofakham, S., and Halassa, M.M. (2017). Thalamic amplification of cortical connectivity sustains attentional control. Nature 545, 219.

Schoenbaum, G., and Eichenbaum, H. (1995). Information coding in the rodent prefrontal cortex. I. Single-neuron activity in orbitofrontal cortex compared with that in pyriform cortex. J. Neurophysiol. 74, 733–750.

Shizuka, D., and McDonald, D.B. (2015). The network motif architecture of dominance hierarchies. J. R. Soc. Interface 12.

Smith, L.B., and Walker, W.H. (2014). The regulation of spermatogenesis by androgens. Semin Cell Dev Biol 30, 2–13.

Smith, J.E., Powning, K.S., Dawes, S.E., Estrada, J.R., Hopper, A.L., Piotrowski, S.L., and Holekamp, K.E. (2011). Greetings promote cooperation and reinforce social bonds among spotted hyaenas. Anim. Behav. 81, 401–415.

Sparta, D.R., Hovelsø, N., Mason, A.O., Kantak, P.A., Ung, R.L., Decot, H.K., and Stuber, G.D. (2014). Activation of Prefrontal Cortical Parvalbumin Interneurons Facilitates Extinction of Reward-Seeking Behavior. J. Neurosci. 34, 3699.

Stockley, P., and Bro-Jørgensen, J. (2011). Female competition and its evolutionary consequences in mammals. Biol. Rev. 86, 341–366.

Taborsky, B., and Oliveira, R.F. (2012). Social competence: an evolutionary approach. Trends Ecol. Evol. 27, 679–688.

Taha, M., McMillon, R., Napier, A., and Wekesa, K.S. (2009). Extracts from salivary glands stimulate aggression and inositol-1, 4, 5-triphosphate (IP3) production in the vomeronasal organ of mice. Physiol. Behav. 98, 147–155.

Takahashi, A., and Miczek, K.A. (2014). Neurogenetics of Aggressive Behavior: Studies in Rodents. In Neuroscience of Aggression, K.A. Miczek, and A. Meyer-Lindenberg, eds. (Berlin, Heidelberg: Springer Berlin Heidelberg), pp. 3–44.

Toyoda, H., Li, X.-Y., Wu, L.-J., Zhao, M.-G., Descalzi, G., Chen, T., Koga, K., and Zhuo, M. (2011). Interplay of Amygdala and Cingulate Plasticity in Emotional Fear. Neural Plast. 2011, 9.

Van Uytfanghe, K., Stöckl, D., Kaufman, J.M., Fiers, T., Ross, H.A., De Leenheer, A.P., and Thienpont, L.M. (2004). Evaluation of a Candidate Reference Measurement Procedure for Serum Free Testosterone Based on Ultrafiltration and Isotope Dilution–Gas Chromatography–Mass Spectrometry. Clin. Chem. 50, 2101.

Vong, L., Ye, C., Yang, Z., Choi, B., Chua Jr, S., and Lowell, B.B. (2011). Leptin Action on GABAergic Neurons Prevents Obesity and Reduces Inhibitory Tone to POMC Neurons. Neuron 71, 142–154.

Wang, F., Zhu, J., Zhu, H., Zhang, Q., Lin, Z., and Hu, H. (2011). Bidirectional Control of Social Hierarchy by Synaptic Efficacy in Medial Prefrontal Cortex. Science 334, 693–697.

Wang, F., Kessels, H.W., and Hu, H. (2014). The mouse that roared: neural mechanisms of social hierarchy. Trends Neurosci 37, 674–682.

Wickersham, I.R., Lyon, D.C., Barnard, R.J.O., Mori, T., Finke, S., Conzelmann, K.-K., Young, J.A.T., and Callaway, E.M. (2007). Monosynaptic Restriction of Transsynaptic Tracing from Single, Genetically Targeted Neurons. Neuron 53, 639–647.

Williamson, C.M., Lee, W., DeCasien, A.R., Lanham, A., Romeo, R.D., and Curley, J.P. (2019). Social hierarchy position in female mice is associated with plasma corticosterone levels and hypothalamic gene expression. Sci. Rep. 9, 7324.

Wilson, E.O. (1975). Sociobiology: The New Synthesis (Cambridge, MA: Havard Publishing).

Wu, Z., Autry, A.E., Bergan, J.F., Watabe-Uchida, M., and Dulac, C.G. (2014). Galanin neurons in the medial preoptic area govern parental behaviour. Nature 509, 325–330.

Xue, M., Atallah, B.V., and Scanziani, M. (2014). Equalizing excitation–inhibition ratios across visual cortical neurons. Nature 511, 596.

Zheng, G.X.Y., Terry, J.M., Belgrader, P., Ryvkin, P., Bent, Z.W., Wilson, R., Ziraldo, S.B., Wheeler, T.D., McDermott, G.P., Zhu, J., et al. (2017). Massively parallel digital transcriptional profiling of single cells. Nat Commun 8, 14049.

Zhou, T., Zhu, H., Fan, Z., Wang, F., Chen, Y., Liang, H., Yang, Z., Zhang, L., Lin, L., Zhan, Y., et al. (2017). History of winning remodels thalamo-PFC circuit to reinforce social dominance. Science 357, 162–168.

Zink, C.F., Tong, Y., Chen, Q., Bassett, D.S., Stein, J.L., and Meyer-Lindenberg, A. (2008). Know Your Place: Neural Processing of Social Hierarchy in Humans. Neuron 58, 273–283.

## Zotero References

Bronson, F.H. (1979). The Reproductive Ecology of the House Mouse. Q. Rev. Biol. 54, 265–299.

Creel, S. (2001). Social dominance and stress hormones. Trends Ecol. Evol. 16, 491–497.

Elo, A.E. (1978). The Rating of Chess Players, Past and Present (New York: Arco).

Gröschl, M. (2009). The physiological role of hormones in saliva. BioEssays 31, 843–852.

Guhl, A.M. (1956). The social order of chickens. Sci. Am. 194, 42–46.

Hille, B. (2001). Ionic Channels of Excitable Membranes (Sinauer Associates, Inc.).

Regehr, W.G. (2012). Short-Term Presynaptic Plasticity. Cold Spring Harb. Perspect. Biol. 4.

Sapolsky, R.M. (2005). The influence of social hierarchy on primate health. Science 308, 648–652.

Wilson, E.O. (1975). Sociobiology: The New Synthesis (Cambridge, MA: Havard Publishing).

